# Sexually dimorphic roles for the type 2 diabetes-associated *C2cd4b* gene in murine glucose homeostasis

**DOI:** 10.1101/2020.05.18.099200

**Authors:** S. Neda Mousavy Gharavy, Bryn Owen, Steven J. Millership, Pauline Chabosseau, Grazia Pizza, Aida Martinez-Sanchez, Emirhan Tasoez, Eleni Georgiadou, Ming Hu, Nicholas H.F. Fine, David A. Jacobson, Matthew Dickerson, Olof Idevall-Hagren, Alex Montoya, Holger Kramer, Zenobia Mehta, Dominic J. Withers, Nikolay Ninov, Paul J. Gadue, Fabian L. Cardenas-Diaz, Céline Cruciani-Guglielmacci, Christophe Magnan, Mark Ibberson, Isabelle Leclerc, Marianne Voz, Guy A. Rutter

## Abstract

Variants close to the *VPS13C/C2CD4A/C2CD4B* locus are associated with altered risk of type 2 diabetes in genome-wide association studies. Whilst previous functional work has suggested roles for *VPS13C* and *C2CD4A* in disease development, none has explored the role of *C2CD4B*. Here, we show that systemic inactivation of *C2cd4b* in mice leads to marked, but highly sexually dimorphic, changes in body weight and glucose homeostasis. Female *C2cd4b* mice display unchanged body weight but abnormal glucose tolerance and defective *in vivo,* but not *in vitro,* insulin secretion, associated with a marked decrease in follicle stimulating hormone levels. In sharp contrast, male *C2cd4b* null mice displayed normal glucose tolerance but an increase in body weight and fasting glycemia after maintenance on high fat diet. No metabolic disturbances were observed after global inactivation of *C2cd4a* in mice, or in pancreatic β cell function at larval stages in *C2cd4ab* null zebrafish. These studies suggest that *C2cd4b* may act centrally to influence sex-dependent circuits which control pancreatic β cell function and glucose tolerance in rodents. However, the absence of sexual dimorphism in the impact of diabetes risk variants argues for additional roles for *C2CD4A* or *VPS13C* in the control of glucose homeostasis in man.

## Introduction

Type 2 diabetes risk is the product of both environmental and genetic factors. More than 200 *loci* have now been described as affecting risk score (1). Whilst most of these affect insulin secretion (2), the identified variants usually lie within or between neighbouring genes and in only a few cases have the causal gene(s) been firmly established (3,4).

Chromosome 15q hosts a risk locus close to the *VPS13C*, *C2CD4A* and *C2CD4B* genes (5) associated with impaired proinsulin processing. Deletion of *Vps13c*, which encodes a lipid transporter (6), selectively from the pancreatic β cell (7), has little effect on glucose homeostasis in the mouse. This finding argues that the latter gene contributes only to a limited extent towards the effect(s) of risk variants in man. Previous expression quantitative trait (eQTL) studies have demonstrated altered expression of *VPS13C* and *C2CD4A* (7), as well as *C2CD4B* (8) in islets from subjects carrying the risk variants. Recently, Kycia *et al.* (9) reported that expression of *C2CD4B*, but not *C2CD4A* or *VPS13C*, was affected by risk alleles. However, the direction of effects of risk alleles differed between the reports, with expression lowered, uniquely in females, in the study of Mehta *et al.* (7) but increased in that of Kycia *et al.* (9). However, both of the latter studies involved relatively small sample numbers. These limitations emphasise the need for interventional studies, involving gene inactivation in tractable model systems such as rodents or fish, as a means of understanding the roles of these genes in metabolic homeostasis.

*C2CD4A* and *C2CD4B* (also called *NLF1* and *NLF2*) (10) encode low molecular mass (39 kDa) proteins of presently unknown function. Unlike the homologous *C2CD4C* gene (expressed from a distinct locus on chromosome 19 in *H. sapiens*), neither C2CD4A nor C2CD4B possess a canonical Ca^2+^/phospholipid binding C2-domain (11). A partly functional Ca^2+^ binding domain may be present on C2CD4B (Supp. Fig.1). Given the essential role for Ca^2+^ in the control of insulin and other hormone secretion (12), an interaction with Ca^2+^ might provide a means through which C2CD4A or C2CD4B influence these processes.

Originally described in endothelial cells as having a largely nuclear distribution, and inducible by cytokines (10), the role of C2CD4B has not been explored previously in β or other metabolically-relevant cell types. Nonetheless, silencing of the single homologous *C2cd4a/b* gene in the zebrafish *(Danio rerio)* led to a decrease in β cell mass (13). In contrast, silencing of the *Drosophila Melanogaster C2CD4A* homologue, *Spenito*, increased circulating levels of the insulin-like molecule IIp2HF (14). Inactivation of *C2cd4c* has no detectable effect on pancreatic development in mice (11).

A recent study (15) has indicated that regulation of *C2cd4a* expression in islets by FOXO1 may be important for the control of insulin secretion. Thus, animals inactivated selectively in the β cell for *C2cd4a* displayed abnormal insulin secretion in response to glucose plus arginine (the effects on secretion stimulated by glucose alone were not reported), and glucose intolerance *in vivo*, and the abnormal expression of β cell signature and “disallowed” genes (16). However, these studies used the *Ins2*-dependent RIP promoter to drive *Cre* expression, a strategy complicated by off-target recombination and ectopic expression of human growth hormone (17).

Here, we use the more direct approach of inactivating *C2cd4b* and *C2cd4a* globally in the mouse, and of deleting its homologue, *C2cd4ab*, in the developing zebrafish (*D. rerio)*, to explore the role of these genes in glucose homeostasis.

## Research Design and Methods

### Animal work

*C2cd4a* (C2cd4a-Del1724-EM1-B6N) and *C2cd4b* (*C2cd4b*^*em2Wtsi*^) mouse strains were generated at the International Mouse Phenotyping Consortium (IMPC), using CRISPR/Cas9. Lean and fat mass were measured using an EchoMRI Quantitative Whole Body Composition analyser (Zinsser Analytic, USA) on unanaesthetised animals.

### Glucose homeostasis

Animals were fasted overnight prior to experiments. For intraperitoneal glucose tolerance tests (IPGTT), glucose (1 g/kg body weight) was injected into the abdomen. In oral glucose tolerance test (OGTT), glucose (2 g/kg body weight) was administered directly into the gut via oral gavage. Blood glucose levels were recorded using an automatic glucometer (Accucheck). For insulin tolerance tests, animals were fasted for 5 h prior to experiments. Insulin was injected into the abdomen (concentrations indicated in the main text). Blood glucose levels were measured post-injection at the time points indicated. To measure insulin secretion *in vivo*, animals were fasted overnight and glucose (3 g/Kg body weight) injected into the abdomen. Blood insulin levels were measured using an Ultra-Sensitive Mouse Insulin ELISA Kit (Crystal Chem, 90080).

### Circulating follicle-stimulating and luteinizing hormone levels

Gonadectomy was conducted under isoflurane anaesthesia, and aseptic conditions, according to standard protocols. Animals received post-operative analgesia, and were allowed to recover for two weeks before collection of blood for hormone analysis (18).

### Insulin secretion from isolated islets

Islets were isolated after pancreatic distension with collagenase, essentially as previously described (19). Insulin secretion was measured as described (20) with 10 size-matched islets in triplicate incubated in Krebs-HEPES-bicarbonate (KREBH) solution (20) containing either: 3 mM glucose, 17 mM glucose or 20 mM KCl at 37°C with gentle shaking. Insulin content was measured using an Insulin Ultra-Sensitive Kit (Supplementary methods).

### Generation of C2CD4A and C2CD4B -FLAG and -GFP tagged constructs

Human *C2CD4A* and *C2CD4B* cDNA sequences were cloned in-frame into plasmid P3XFLAG-CMV-14 (Addgene) to provide a C-terminal 3xFLAG epitope tag. Green fluorescent protein (GFP)-tagged proteins were generated by inserting the human *C2CD4A* and *-B* cDNA sequences into the C-terminus of GFP using plasmid pEGFP-C1 (Addgene and Clontech).

### Time-lapse imaging

Cells were grown on coverslips and transfected with either GFP-tagged C2CD4A, C2CD4B or Syt1-containing constructs. 24 h post-transfection, cells were incubated for 1 h at 37°C with aerated KREBH solution.

### Statistical analysis

Data were analysed using GraphPad prism 8.0. p-values <0.05 were considered significant.

### Study approval

All mouse *in vivo* procedures were conducted in accordance with the UK Home Office Animal (Scientific Procedures) Act of 1986 (Project license PA03F7F0F to I.L.) as well as being approved by Imperial College Animal Welfare and Ethical Review Body. All zebrafish work was approved by the ethical committee of the University of Liège (protocol number 13-1557) or under European Union and German laws (Tierschutzgesetz), and with the approval of the TU Dresden and the Landesdirektion Sachsen (approval license number: TVV 45/2018).

Additional methods are included in a separate file (Supplementary Methods).

## Results

### *C2CD4A* and *C2CD4B* expression in mouse and human islets

Human C2CD4A and C2CD4B are 83 % homologous. Like their murine homologues, the two human genes are predicted to have evolved from a common ancestor (see phylogenetic tree, Supp. Fig.2, A). Conservation of genomic architecture (synteny) at this locus argues for direct homology between the human and murine forms of each gene. The zebrafish *D. rerio* possesses two *C2cd4*-like genes homologous to *H. sapiens* and *M. musculus*: *C2cd4a* and *C2cd4c* (Supp. Fig.2, A). Analysis of gene expression databases (Biogps.org.), and previous publications (21) (22), reveals ~10-fold higher levels of *C2cd4b* than *C2cd4a* mRNA in mouse islets and purified mouse β cells (Supp. Table 1). In contrast, roughly equal levels of *C2CD4A* and *C2CD4B* mRNA are present in human islets (23) and purified β cells (24). Expression of both genes was detected in the pituitary in both human and mouse (Data Source: GTEx data, analysed in The Human Protein Atlas, URL: https://www.proteinatlas.org/ENSG00000198535-C2CD4A/tissue, https://www.proteinatlas.org/ENSG00000205502-C2CD4B/tissue). *C2cd4a* and *C2cd4b* were both upregulated in pancreatic islets of the high fat diet-fed DBA2J diabetic mouse model (Supp. Fig.3,A,B) (25). No evidence of *C2CD4A or C2CD4B* upregulation was observed in the islets of patients with T2D *versus* normoglycemic controls (26).

Examination of the human *VPS13C/C2CD4A/C2CD4B* locus revealed multiple regulatory elements (Fig.1), consistent with recent findings (9). A single nucleotide polymorphism, rs7163757, has recently been shown by fine mapping as the likely causal variant for T2D risk at this locus (9). Correspondingly, CRISPRa or CRISPRi at the rs7163757 site has the most marked effects on neighbouring genes (27), consistent with this being the effector variant (SNP). To explore in more detail the potential role of the enhancer around the diabetes risk variant rs7163757 (9), and whether it may play a role in gene expression in disease-relevant tissues (notably the islet and brain/pituitary), we introduced a reporter bearing 1303 nucleotides of the human sequence into the zebrafish genome, controlling the production of GFP from a minimal c*Fos* promoter (Supp. Fig.2, B). Expression was restricted to the endocrine pancreas and brain at all stages (Supp. Fig. 2, C), and was detected in all islet cell types, most strongly in delta cells (Supp. Fig.2, D).

**Figure 1.**
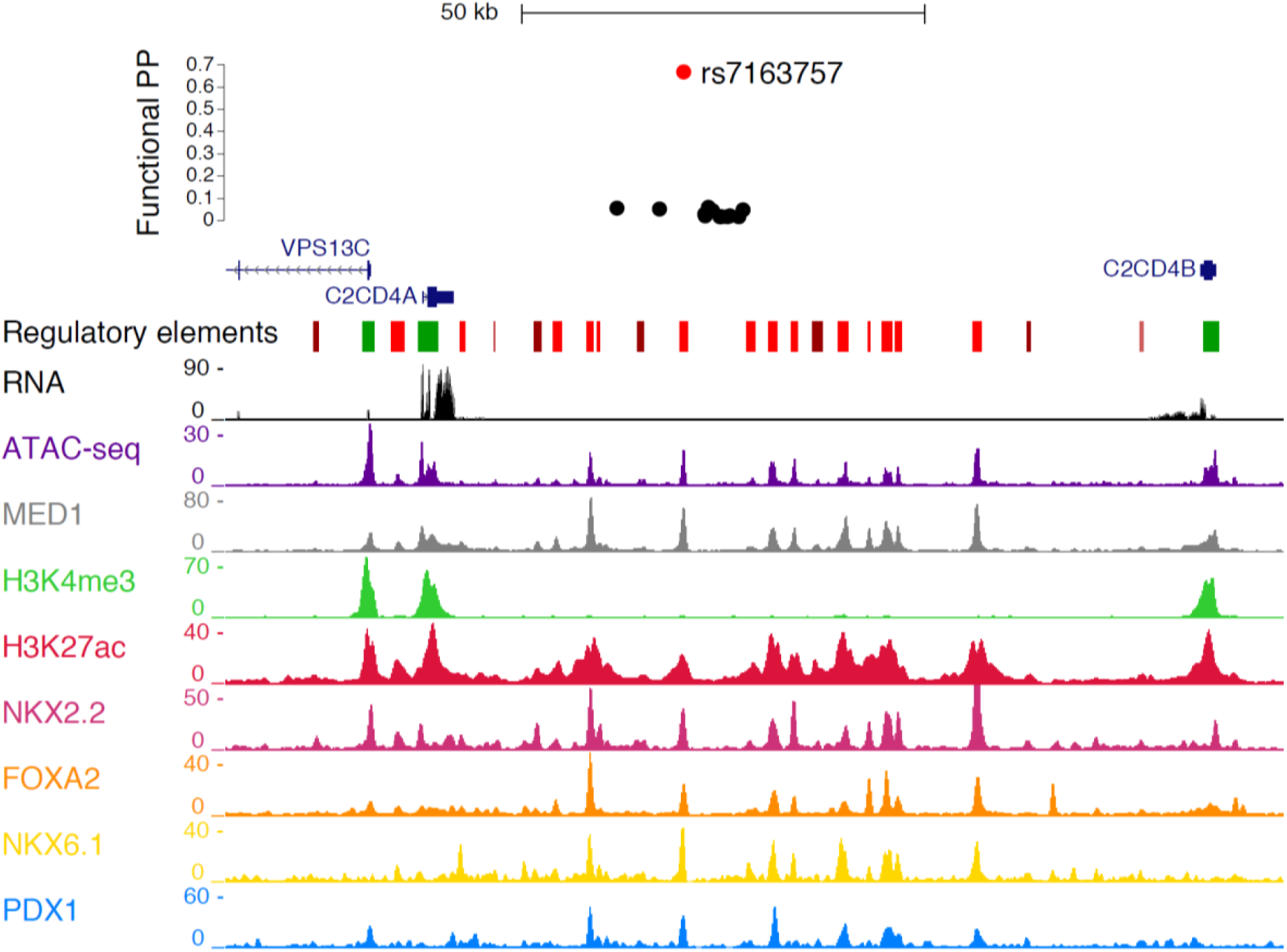
Genomic context of T2D variants in the *VPS13C/C2CD4A/C2CD4B* locus. SNP at rs7163757 is located in an open chromatin region between *C2CD4A* and *C2CD4B*, as assessed by Assay for Transposase-Accessible Chromatin sequencing (ATAC-seq) data. Chromatin immunoprecipitation and next generation sequencing (ChIP-seq) data reveals binding sites for transcription factors involved in the development and function of β cells, including FOXA2, NKX2.2, NKX6.1 and PDX1. Data from Miguel-Escalada *et al*., *Nature Genetics* 2018 (27).

### Role of *c2cd4ab* in the zebrafish larvae

Zebrafish possess a single gene, *c2cd4ab*, homologous to the two mammalian counterparts (Supp. Fig.2, A). As a convenient proxy for insulin secretion in the larvae of fish inactivated for *c2cd4ab* or in wild-type controls (see Methods), glucose-stimulated Ca^2+^ dynamics were monitored *in vivo* by imaging the fluorescence of a gCaM6 transgene. Ca^2+^ changes did not differ between wild type and mutant animals (Fig.2, A-E), arguing against a role for *c2cd4ab* in β cell function during development.

**Figure 2.**
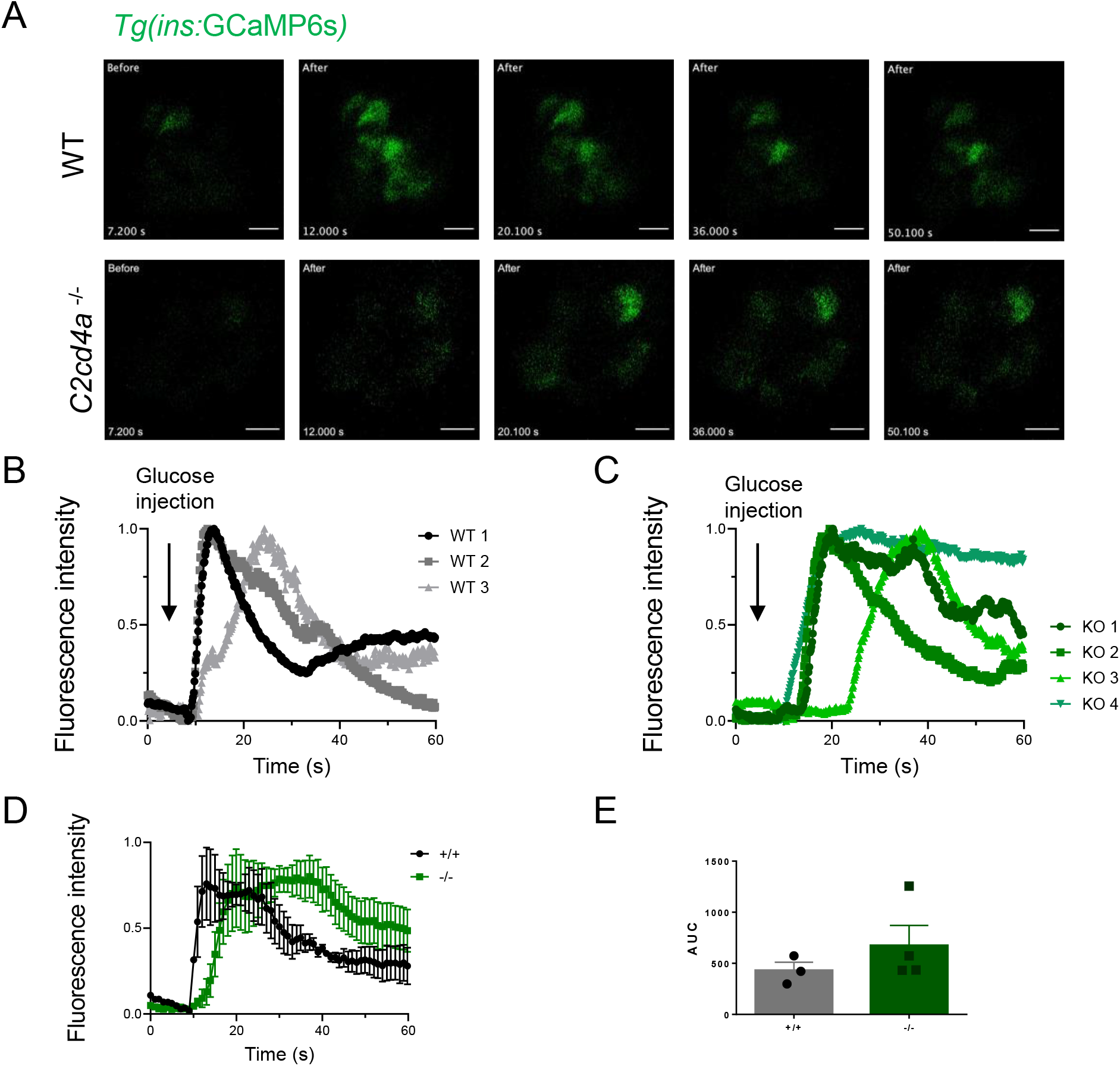
Glucose-stimulated calcium influx is not altered by *c2cd4a* deficiency in zebrafish larvae. A. Representative time series showing single confocal planes of islets in WT and *c2cd4a-/-* zebrafish larvae (upper and lower panel respectively) before and after pericardial injection of 25 mM glucose. The larvae express the genetically encoded Ca^2+^ indicator GCaMP6s (green) in their β cells, which exhibits an increase in fluorescence upon glucose stimulation in both WT and *C2cd4a*^*−/−*^ larvae. Glucose injection took place after 7 sec of imaging. B-D. Plots showing normalized islet GCaMP6s fluorescence intensity: (F_t_ - F_Min_)/(F_max_ - F_min_), over time for different islets from WT (B) (WT, n=3), *C2cd4a*^*−/−*^ larvae (C) (*C2cd4a*^*−/−*^, n=4) and the averages of both (D) with area under the curve quantified in E. The arrows indicate the time of glucose injection, showing a consistent increase in GCaMP6s fluorescence upon glucose injection in WT and *C2cd4a*^*−/−*^ larvae. Scale bars= 10 μm.

### Female *C2cd4b* null mice are glucose intolerant and display defective insulin secretion *in vivo*

Given the substantially higher expression of *C2cd4b* than *C2cd4a* in mouse islets (Supp. Table 1), we studied mice in which the former gene was deleted globally (International mouse phenotyping consortium, IMPC, https://www.mousephenotype.org/) (Fig.3, A). Inter-crossing of heterozygous animals produced pups at the expected Mendelian ratio (Fig.3, B) and resulted in complete elimination of *C2cd4b* mRNA from isolated islets (Fig.3, C). Female *C2cd4b* null mice gained weight at the same rate as control littermates whether maintained on regular chow (RC) or a high -fat and -sucrose diet (HFD) (Fig.3, D), and no differences were apparent in fed or fasting glycemia either on RC (Supp. Fig.4, A,C) or HFD (Fig.3, F and Supp. Fig.4, E). Intraperitoneal glucose tolerance was examined for animals maintained on RC (Supp. Fig.5) or HFD (Supp. Fig.6) from 8-22 weeks of age. Females displayed abnormalities from 12 weeks of age on RC (Supp. Fig.5, A,C,E,G), and 8 weeks on HFD (Supp. Fig.6, A,C,E) with data at 22 weeks shown in Fig.4, A,C. Whereas oral glucose tolerance was normal on RC (not shown), defects were observed on HFD (Fig.4, E). For female mice on RC, these changes were associated with defective insulin secretion *in vivo* (Fig.5, A) as also indicated by unchanged insulin levels after glucose injection despite elevated plasma glucose levels in female knockout mice maintained on HFD (Fig.5, C). This was despite any reduction in β cell mass, as assessed by histochemical analysis of pancreatic slices (Supp. Fig.7, A-D). Furthermore, we observed no differences in *C2cd4b* null female mice by homeostasis model assessment of β cell function (HOMA2-%B) (Supp. Fig.8, A,C). Likewise, insulin sensitivity was not different between *C2cd4b* null and wild type mice (Supp. Fig.9, A,C).

**Figure 3.**
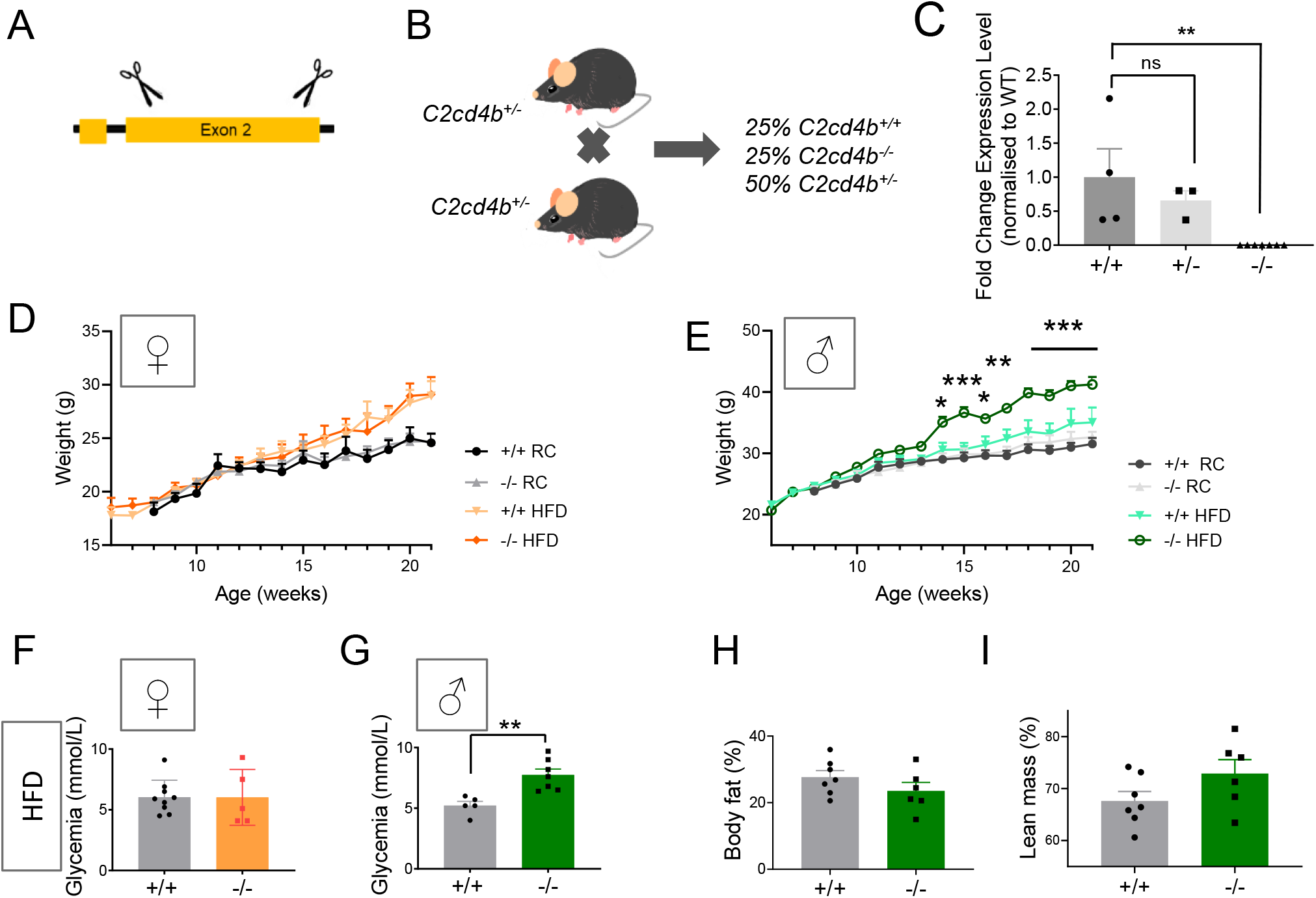
Characterisation of *C2cd4b* null mice. A. *C2cd4b* global null mice (*C2cd4b*^*em2Wtsi*^) were generated by the International Mouse Phenotyping Consortium (IMPC). Using CRISPR/Cas9, the encoding exon from murine *C2cd4b* (exon 2) was deleted. B. *C2cd4b*^*+/−*^ (heterozygous) animals were setup as breeding pairs and the wild type (*C2cd4b*^*+/+*^), and homozygous (*C2cd4b*^*−/−*^) littermates were studied. C. RT-q-PCR performed on RNA from isolated islets showed a significant decrease in *C2cd4b* mRNA levels in homozygous animals (p=0.0092), (*C2cd4b*^*+/+*^ n=4, *C2cd4b*^*+/−*^ n=3, *C2cd4b*^*−/−*^ n=7). **p<0.01, data were assessed for significance using an unpaired Student’s t-test. D-E. Changes in weight of *C2cd4b*^*+/+*^ and *C2cd4b*^*−/−*^ female (D) and male (E) mice over time on regular chow diet (RC) or high-fat and -sucrose diet (HFD), (number of animals assessed on RC: F^*+/+*^ n=5-6 and F^*−/−*^n=7-13; M^*+/+*^ n=7-12 and M^*−/−*^n=10-12. HFD: F^*+/+*^ n=10-12 and M^*−/−*^n=7-10; M^*+/+*^ n=8-12 and M^*−/−*^n=10-12). F-G. Fasting glycemia in animals on HFD were measured at 23 weeks of age (number of animals used : F^*+/+*^ n=9, F^*−/−*^n=5; M^*+/+*^ n=5, M^*−/−*^n=7). H-I. Percentage of body fat (H) and lean mass (I) in males maintained on HFD at 20 weeks of age. *\**p<0.05, **p<0.01, ***p<0.001, the results were assessed for significance using a mixed-effect analysis, Tukey’s multiple comparison test. Data were assessed for significance using an unpaired Student’s t-test, or 2-way ANOVA where two genotypes were compared. Values represent means ± SEM.

**Figure 4.**
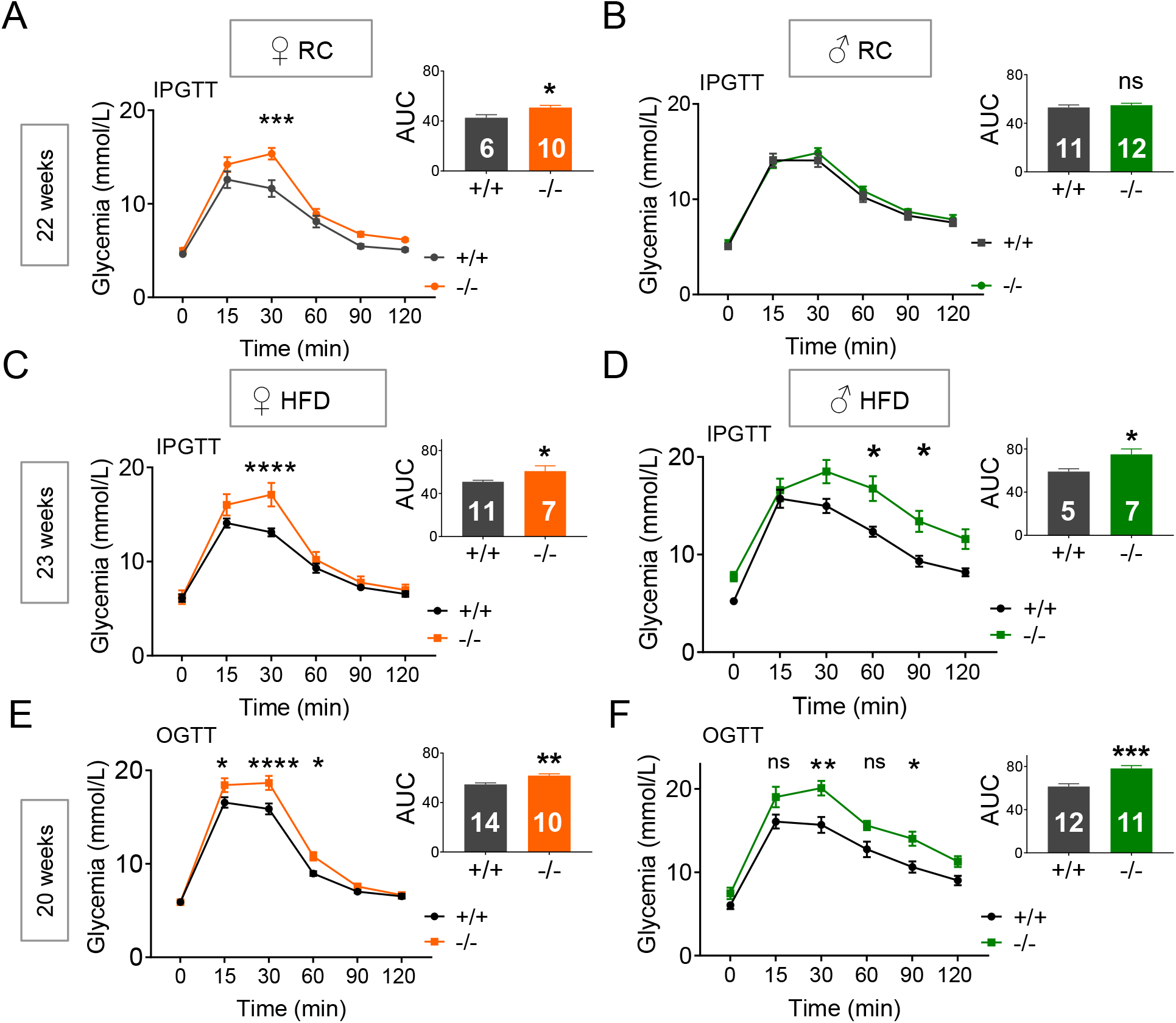
*C2cd4b* null mice display glucose intolerance in glucose tolerance tests (IPGTTs/OGTTs). A-B. IPGTTs were performed on female (A) and male (B) mice at 22 weeks of age maintained on RC. C-D. IPGTTs were performed on *C2cd4b* female (C) and male (D) mice at 23 weeks of age maintained on HFD. E-F. Oral glucose tolerance tests (OGTTs) were performed on *C2cd4b* female (E) and male (F) mice at 20 weeks of age. *p<0.05, **p<0.01, ***p<0.001, data were assessed for significance using a 2-way ANOVA with Bonferroni’s multiple comparison test. Inset: area under the curve (AUC) analysis; assessed for significance using an unpaired Student’s t-test. Numbers in bar graphs represent the number of animals used (same number of samples used for glycemia and AUC graphs). Values represent means ± SEM.

**Figure 5.**
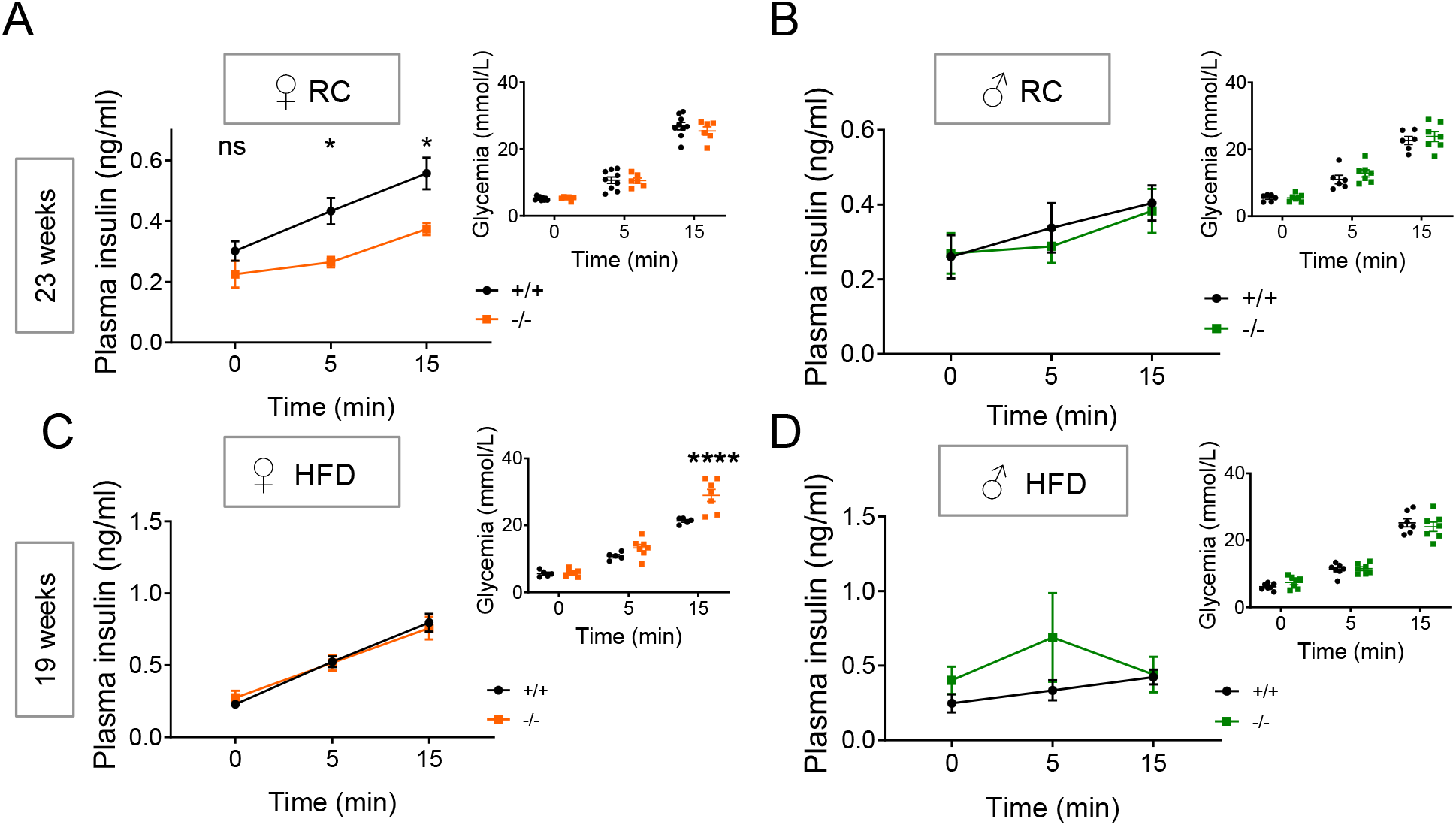
Effect of *C2cd4b* deletion on *in vivo* glucose-stimulated insulin secretion. *In vivo* insulin levels during IPGTT in *C2cd4b* mice on RC (A, B) and HFD (C, D). (RC: F^*+/+*^ n=9, F^*−/−*^n=6, M^*+/+*^ n=7, M^*−/−*^n=8; HFD: F^*+/+*^ n=5 F^*−/−*^n=7, M n= 6). * p<0.05, **p<0.01, ***p<0.001, data were assessed for significance using a 2-way ANOVA with Bonferroni’s multiple comparison test. Values represent mean ± SEM.

Examined *in vitro*, glucose-stimulated insulin secretion was not different between islets from wild type or *C2cd4b* null female mice maintained on RC diet (Supp Fig. 10, A), but was slightly elevated in those from female null mice maintained on HFD (Supp. Fig.10, C). Likewise, we observed no alterations in glucose or KCl-stimulated stimulated Ca^2+^ dynamics (Supp. Fig.11, A,B) or in β cell-β cell coupling (Supp. Fig.11, C-F). Correspondingly, no changes in voltage-activated Ca^2+^ currents were apparent in patch clap recordings (Supp. Fig.12, A-C). Massive parallel sequencing (RNA-Seq; Supp. Table 2) of islets from *C2cd4b* null or control animals confirmed the lowering of *C2cd4b* expression, and revealed a significant (~75%) increase in *C2cd4a* expression. No other mRNAs were significantly affected by *C2cd4b* deletion (Supp. Table 2).

Given the absence of clear insulin secretion defects in isolated *C2cd4b* null islets, and the expression of both *C2cd4a* and *C2cd4b* in the pituitary (see above), we next assessed whether deletion of this gene might affect the production of sex hormones and thus provoke gender-specific differences in glucose homeostasis. In order to remove a negative feedback loop through which hormones secreted from the gonads repress follicle-stimulating hormone (FSH) and luteinizing hormone (LH) release from the pituitary gland, animals were gonadectomised prior to these experiments. Compared to wild type littermates, female *C2cd4b* null mice displayed ~ 50 % lower circulating FSH levels (Fig.6, A,C). In contrast, no differences in LH or FSH levels were apparent between wild type and *C2cd4b* null male mice (Fig.6, B,D).

**Figure 6.**
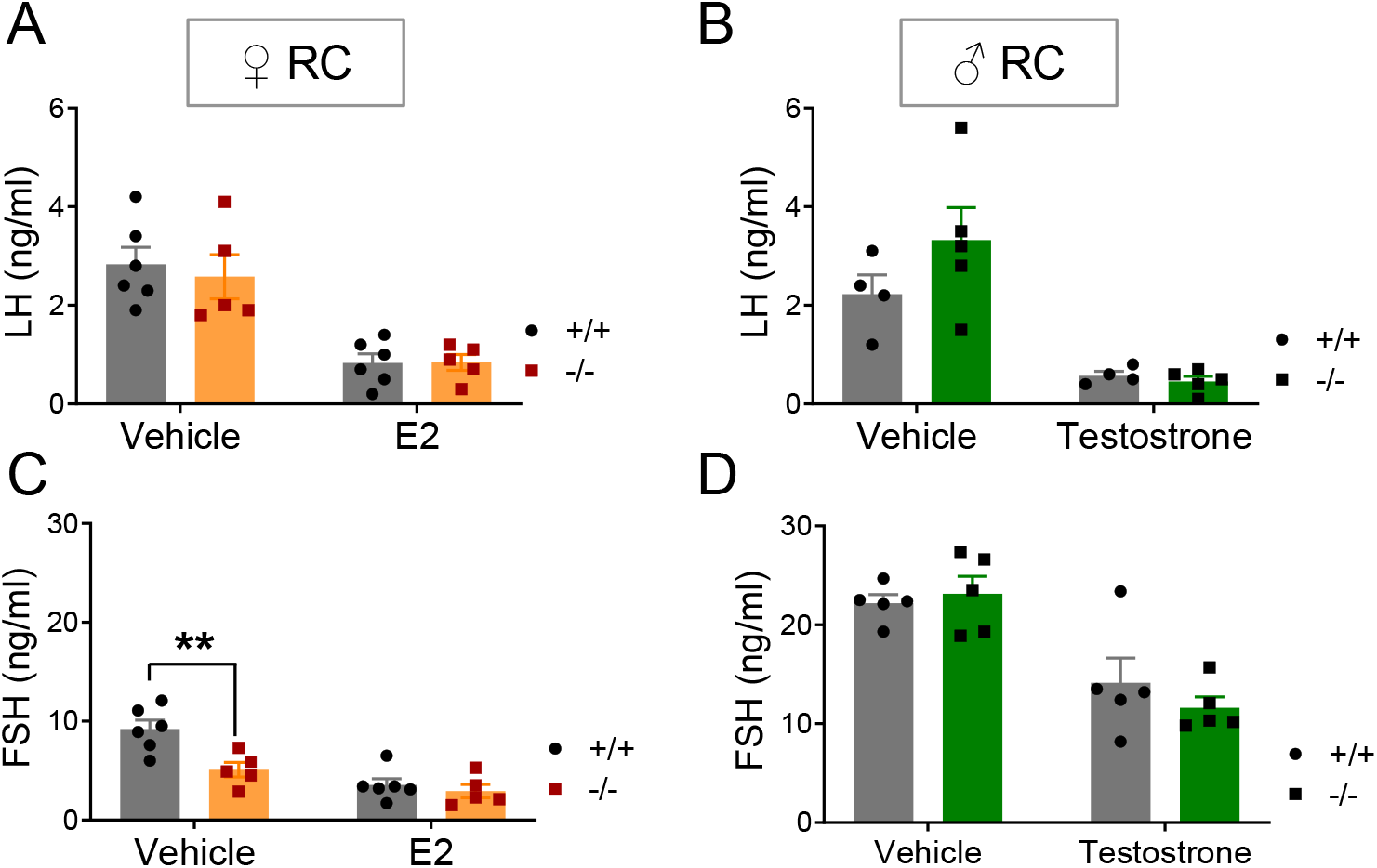
Effect of deletion of *C2cd4b* on FSH and LH release from the pituitary gland on gonadectomised mice. A-B. Circulating LH levels in *C2cd4b* female and male mice after injections with saline (vehicle), estradiol (E2) or testosterone respectively. C-D. Circulating FSH levels in *C2cd4b* female and male mice after injections with saline (vehicle), E2 or testosterone respectively. * p<0.05, **p<0.01, ***p<0.001. Assessed for significance using an unpaired Student’s t-test. n=5 mice/genotype. Values represent mean ± SEM.

### Male *C2cd4b* null mice display elevated body mass and fasting glycemia on a high-fat and -sucrose diet

Maintained on RC diet, male *C2cd4b* null mice gained weight at the same rate as wild type littermates while, in contrast to females, male mutant mice gained substantially more weight on HFD from 14 weeks of age *versus* wild type littermates (Fig.3, E) associated with raised fasting glycaemia (Fig.3,G). This increase in body weight was largely the result of augmented lean mass (Fig.3, H,I). Intraperitoneal glucose tolerance was normal in males maintained on a RC diet at all ages examined (Fig.4, B, Supp. Fig.5, B,D,F,H). Although unaltered in younger male mice after maintenance on HFD (Supp. Fig.6, B,D,F,G), glucose tolerance was impaired at 23 weeks by *C2cd4b* deletion (Fig.4, D,F). As observed in female *C2cd4b* null mice, β cell mass (Supp. Fig.7, E-H), HOMA2-%B (Supp. Fig.8, B,D) insulin sensitivity (Supp. Fig.9, B,D), glucose or KCl-stimulated insulin secretion from isolated islets (Supp. Fig.10, B,D) were unaltered in male *C2cd4b* null mice versus littermate controls.

### *C2cd4a* null mice display no metabolic abnormalities

To determine whether *C2cd4a* inactivation might also impact glucose homeostasis, we next examined male or female *C2cd4a* null mice (Supp. Fig.13, A-C). In contrast to their *C2cd4b* null counterparts, *C2cd4a* null mice displayed no evident metabolic phenotype up to 22 weeks of age and neither weight gain, glucose homeostasis nor insulin secretion differed between wild type and null mice (Supp. Fig.13, D-I) for either sex. Likewise, measured *in vitro*, glucose or KCl (depolarisation)-stimulated insulin secretion were unaltered in isolated islets from female *C2cd4a* null mice (Supp. Fig.13, J).

### C2CD4A but not C2CD4B C2 domains support Ca^2+^-dependent intracellular translocation

We next sought to explore the mechanism(s) through which C2CD4B or C2CD4A may influence β cell (and, potentially, pituitary gonadotroph) function. Both proteins have been suggested (11) to lack a functional C2 domain, consistent with a reported localisation in the nucleus in COS7 cells (10). In contrast to earlier findings reporting nuclear subcellular localisation, when overexpressed as GFP- or FLAG-tagged chimaeras in rodent (MIN6, INS1 (832/13)) or human (EndoCβH1) β cell lines, C2CD4A and C2CD4B were found at the plasma membrane, the cytosol and the nucleus (Supp. Fig.14, A-C and Supp. Fig.15, A,B). In the majority of the cells, C2CD4A and C2CD4B were localised to the cytoplasm and nucleus. Co-localisation with readily-identified intracellular sub-compartments including the secretory granule (insulin), trans-Golgi network (TGN46), endosome/lysosome (LAMP1) or ER (KDEL) was not apparent (Supp. Fig.16, A,B).

The above findings suggested that the C2 domain of either protein may bind to Ca^2+^ and contribute to localisation at, and/or shuttling between, subcellular compartments in living cells. To test this hypothesis, we explored phospholipid-dependent recruitment to the plasma membrane in INS1(832/13) β cells (28) expressing either a control construct, in which Synaptotagmin-1 (Syt1), bearing five C2 domains, was fused to GFP (29), or equivalent C2CD4A or -B constructs (N-terminal linkage). In response to an increase in intracellular free Ca^2+^ provoked by 50 μM extracellular Ca^2+^ and the calcium ionophore, ionomycin (50 ng/ml), Syt1-GFP translocated from intracellular (likely ER-bound) sites to the plasma membrane. This movement was readily visualized by simultaneous live cell wide-field and total internal reflection of fluorescence (TIRF) imaging (Fig.7, A,B,D). A similar, but smaller change in the localisation of C2CD4A-GFP in response to Ca^2+^ was also observed. In contrast, no response was detected for C2CD4B-GFP (Fig.7, C,E-G).

**Figure 7.**
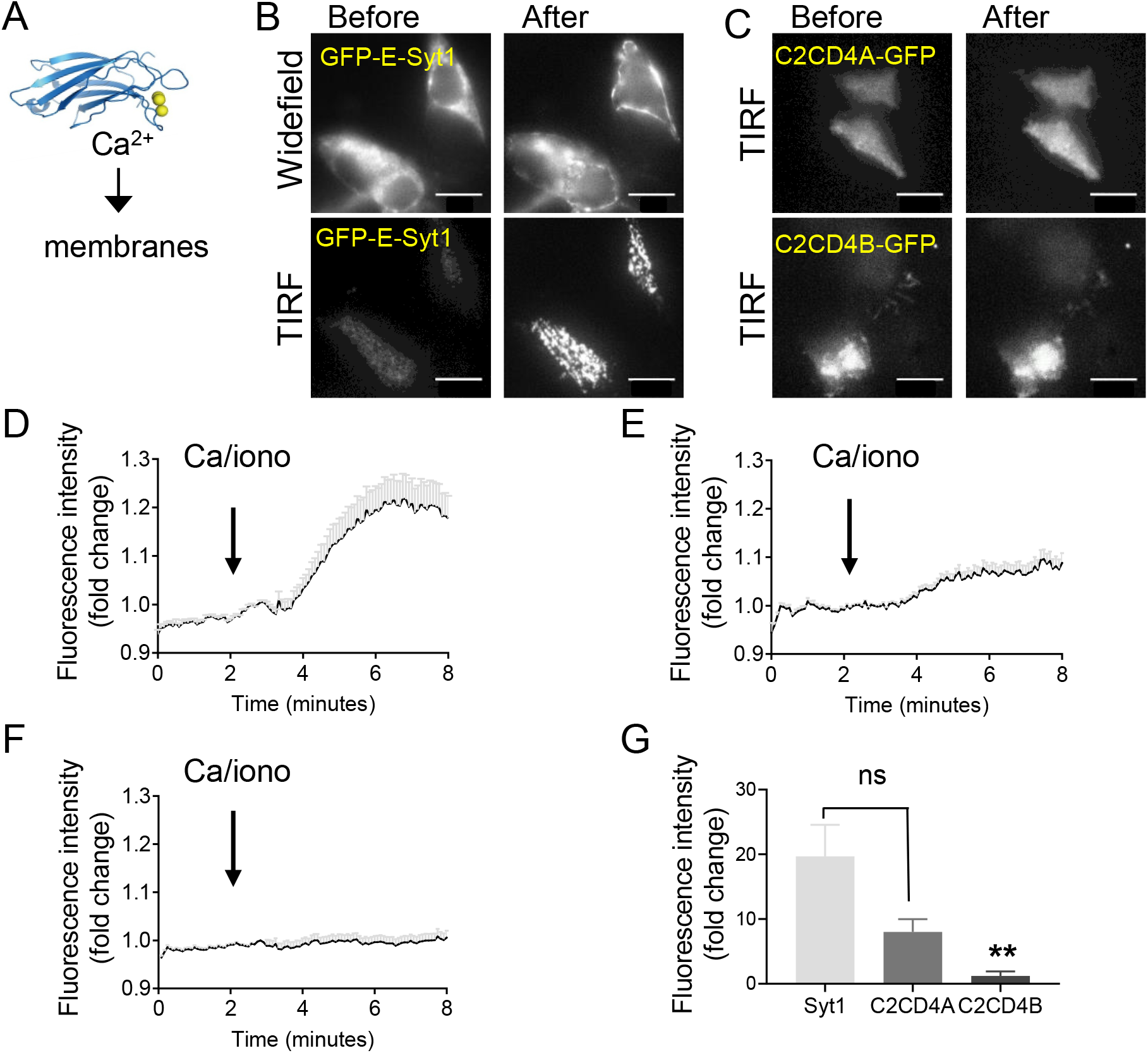
Changes in C2CD4A and C2CD4B localisation in response to increased intracellular [Ca^2+^]. A. Synaptotagmin (Syt1) moves from the ER to the plasma membrane in response to elevated Ca^2+^ (positive control). B. Localisation of GFP-Syt1 at 5 mM glucose shown before and after an increase in intracellular Ca^2+^, achieved by addition of 50 ng/ml ionomycin, obtained by simultaneous wide-field and TIRF image acquisition. C. Localisation of GFP-tagged-C2CD4A and -C2CD4B before and after addition of ionomycin. D-F. Time courses for the translocation of the GFP tagged proteins (Syt1 (D), C2CD4A (E) or C2CD4B (F)) obtained by TIRF imaging before and after addition of ionomycin. G. Assessment of fluorescence intensity fold-change (%) reveals an increase in C2CD4A intensity at the plasma membrane, after the imposed increase in intracellular Ca^2+^ levels, which was significantly higher than that observed for C2CD4B, but not C2CD4A compared to Syt1 protein translocation. *p<0.05, **p<0.01, data were assessed for significance using an ordinary one-way ANOVA. Solid lines represent mean ± SEM; SEM shown by grey bars. Scale bars= 10 μm.

### Identification of C2CD4A and C2CD4B binding partners by mass spectrometry

The above experiments demonstrated that C2CD4A, and possibly C2CD4B, may participate in Ca^2+^-dependent signal transduction. To gain further insight into possible mechanisms of action we performed an unbiased proteomic screen using immunoprecipitation and mass spectrometry to identify potential binding partners. Normalising to the negative control, and ranking in order of protein abundance, we generated lists of possible interacting proteins for human C2CD4A, C2CD4B (Supp. Tables 3,4), or both (Table 1). MIN6 cells transfected with human C2CD4A or C2CD4B were used given the low transfection efficiency of human-derived EndoCβH1 cells (see Supplementary Methods). Interacting partners included proteins involved in Ca^2+^ binding (TOR2A and EFCAB5 (30) and (https://www.genecards.org/), NF-κB signalling (SQSTM1 and PDCD11) and protein trafficking (PCSK9, NEDD4, RAP1 and GAP2). Ptprn (IA-2) and Ptprn2 (phogrin/IA-2β/ICA512) bound to both C2CD4A and C2CD4B. These protein tyrosine phosphatase-like transmembrane proteins are granule-resident, and implicated in granule trafficking and exocytosis (31).

**Table 1.**
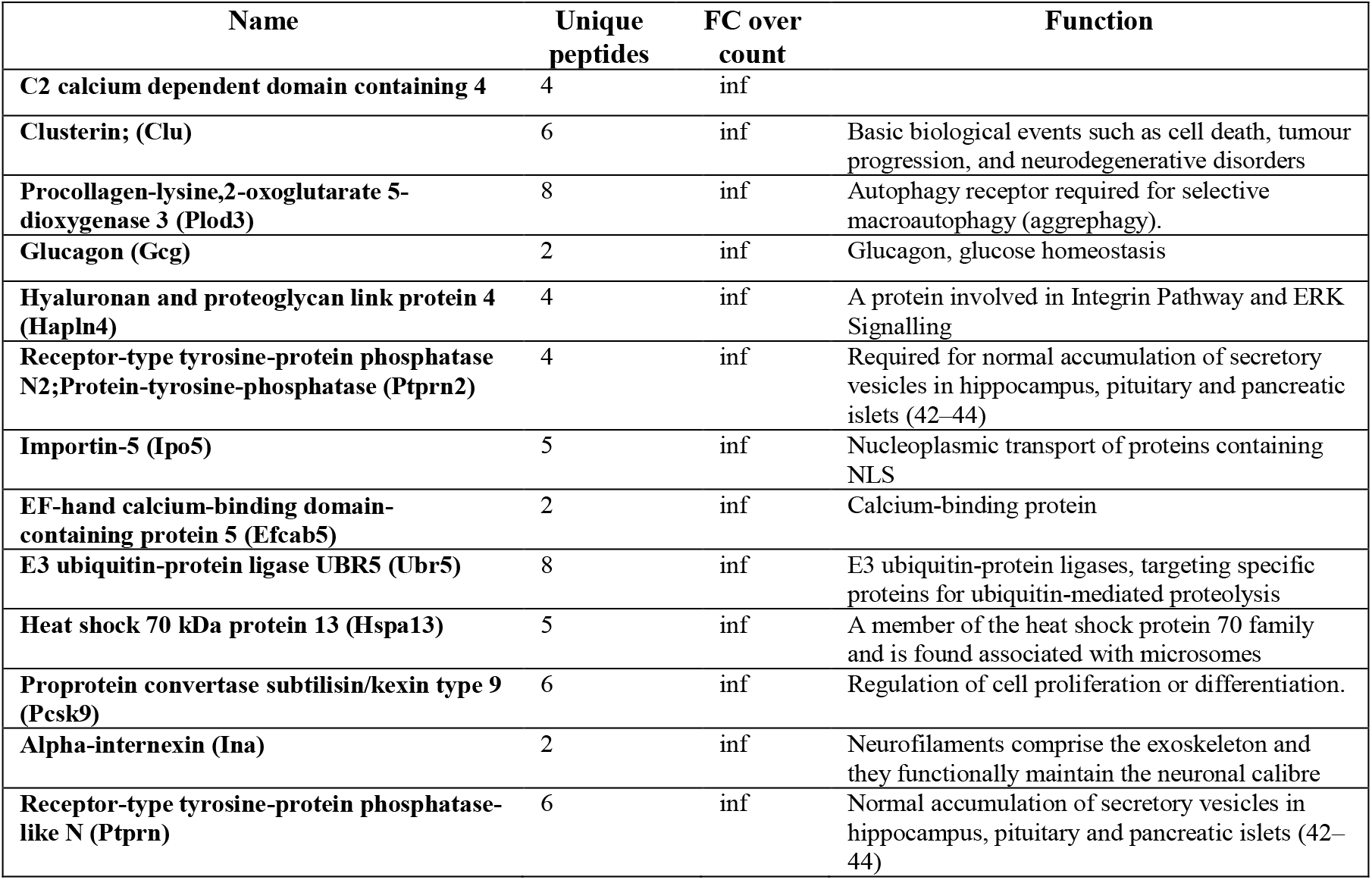
C2CD4A and C2CD4B interacting proteins detected by mass-spectrometry. Top 12 predicted interacting proteins with both C2CD4A and C2CD4B in MIN6 cells. The given function of the identified protein was adopted from GeneCards.org.

## Discussion

The overall aim of this study was to examine the biological roles of *C2cd4b* and *C2cd4a in vivo* focussing on the pancreatic β cell and pituitary. These questions have been pursued using mouse and fish knockout models and relevant cell lines. In contrast to earlier findings (15), we observed that global *C2cd4a* deletion in the mouse exerted no effects on insulin secretion *in vitro* or *in vivo*. These findings are consistent with the considerably (10-fold) lower expression of *C2cd4a* than *C2cd4b* in mouse islets and derived β cell lines (Supp. Table 1) though we would emphasise that no attempt was made here to quantify protein (rather than mRNA) levels. The reasons for the differences between the present study and that of Kuo *et al.* (15) with respect to *C2cd4a* are presently unclear, although this may reflect differences in the targeting strategies used to generate the different mouse mutants or genetic background and general housing conditions. Nonetheless, given that our work involved global inactivation of this gene we cannot exclude the possibility that compensatory changes occur in other tissues, or in islet cell other than β cells, that mitigate the effects of deleting *C2cd4a* in the β cell. Of note, we observed no changes in the expression of “disallowed” genes including *Ldha* after *C2cd4b* deletion, nor in other glycolytic genes such as *AldoA*, as reported after over-expression of *C2cd4a* (15), suggesting non-redundant roles of *C2cd4a* and *C2cd4b*.

### Deletion of *C2cd4b* leads to weight gain in males, but defective insulin secretion in female mice

Our findings reveal a strikingly sexually dimorphic phenotype of *C2cd4b* null mice, which is also strongly dependent upon diet. In contrast to females, male *C2cd4b* null mice displayed no evident metabolic abnormalities at any age when maintained on RC. Although mutant male mice exhibited a substantial increase in body weight on HFD, fasting glucose and glucose tolerance were impaired only in older (>20 weeks) animals. In stark contrast, female mutant mice displayed abnormal glucose tolerance from as early as 12 weeks on both RC and HFD (with a tendency towards abnormal glucose tolerance observed at 8 weeks), despite unaltered body weight. The changes in lean mass observed in *C2cd4b* null mice on HFD are reminiscent of findings from the IMPC (https://www.mousephenotype.org/data/genes/MGI:1922947) when this line was fed RC. Interestingly, these earlier, less in-depth studies, also reported gender-specific alterations in lipid, circulating cholesterol and triglyceride levels in male *C2cd4b* null mice, but failed to detect alterations in glucose tolerance. No metabolic phenotype was reported for *C2cd4a* mutants: (https://www.mousephenotype.org/data/genes/MGI:3645763). We note that a statistically significant (p=0.004) impact of the human T2D-associated variant rs7172432 (which is in perfect linkage disequilibrium with rs7163757: R^2^=1.0) (15) at the *VPS13C/C2CD4A/C2CD4B* locus on waist circumference has previously been reported (32), and may be related to the alterations in body weight we observed here in male *C2cd4b* knockout mice. However, the human studies were not stratified by sex.

What factors underlie these gender-dependent differences in insulin secretion between wild type and *C2cd4b* null mice? Our findings suggest that β cell-extrinsic mechanisms, possibly involving circulating factors, contribute to (or may be the drivers of) altered insulin secretion in the living animal. Potential contributors are changes in FSH levels (33,34) in female null *C2cd4b* mice, reflecting altered expression of the gene in the pituitary. Decreased FSH production is expected, in turn, to decrease circulating oestrogen levels. We note, however, that measurements of estrogen are complicated in fertile mice due to fluctuations in the estrous cycle. Positive actions of estrogen on β cell insulin content (via ERa) and secretion (via ERbeta) are well known (33,35), and may thus underlie the weaker insulin secretion in *C2cd4b* knockout mice. Consistent with a requirement for sexual maturity, no differences were apparent in β cell function between genotypes in the living fish embryo (Fig.2), at stages where differences in circulating sex hormones are not anticipated. Nevertheless, earlier studies in zebrafish larvae using oligonucleotide-mediated *C2cd4ab* gene knockdown (13) reported alterations in β cell mass.

Such sexual dimorphism on the impact of variants at this locus, however, has not been reported in human GWAS data (36,37). One possible explanation is that dimorphism in mice reflects well-known differences in the response of males and females to HFD (38). More likely, in our view, is that in rodent islets (and pituitary; see below) *C2cd4b* expression predominates over *C2cd4a*, while in humans, *C2CD4A* and *C2CD4B* expression are comparable. Consequently, in humans, changes in both *C2CD4A* and *C2CD4B* may mediate the effect of the GWAS signal for T2D, dampening a sexually dimorphic effect of changes in *C2CD4B* expression.

Might altered genetic risk for T2D in man result from changes in *C2CD4A* or *-B* expression in the brain? Importantly, high levels of expression of C2CD4A and -B in pituitary (GTEx data, The Human Protein Atlas, URL: https://www.proteinatlas.org/ENSG00000198535-C2CD4A/tissue, https://www.proteinatlas.org/ENSG00000205502-C2CD4B/tissue) are consistent with this view. Correspondingly, an eQTL for *C2CD4A* is reported for rs7163757 in the pituitary (https://www.gtexportal.org/home/snp/rs7163757#sqtl-block) with a tendency towards decreased expression of *C2CD4A*, and more strongly *VPS13C*, in carriers of C (risk allele) *vs* T alleles.

### Intracellular signalling by *C2cd4a* and *C2cd4b*

Our observation that neither C2CD4A nor C2CD4B are localised exclusively in the nucleus, as previously reported in COS7 cells (10), islets and MIN6 cells (15) was unexpected, but implies a more dynamic role for both C2CD4A, and possibly C2CD4B, in intracellular signalling. We note that the present studies explored the subcellular distribution of the *H. sapiens* protein, rather than the *M. musculus* homologue examined previously (15), providing a potential explanation for these differences. Interestingly, predictions from primary structure (11) indicate that neither C2CD4A nor C2CD4B (human or mouse) possesses a C2 domain with a *bona fide* Ca^2+^ binding site (39,40). Nevertheless, we provide a direct demonstration of Ca^2+^-dependent recruitment to the plasma membrane of C2CD4A (Fig.7) whereas C2CD4B would appear to exert its function independently of calcium binding.

What signalling mechanism(s) may lie downstream of C2CD4A (or C2CD4B) after recruitment to the plasma membrane? Of those proteins identified as interacting with both, Ptprn2 (also known as phogrin and IA-2β) is of particular interest, as it is known to play a role in the control of secretion from neuroendocrine cells (31). Thus, inactivation of Ptprn2/phogrin, a secretory granule-localised protein (41), leads to defective insulin secretion in mice (42,43). Importantly, double knockout of Ptprn2/phogrin and the closely-related *Ptprn (IA-2)* gene leads to defective FSH and LH production and female infertility (42), implying an important role in the anterior pituitary. An interaction between C2CD4A or C2CD4B and PTPRN2 in the pituitary (and possibly the β cell) may therefore contribute to the effects of altered expression on diabetes risk. Whether similar interactions are relevant elsewhere in the brain such as in feeding centres, and contribute to weight gain in male *C2cd4b* null mice, is unknown.

## Conclusions

The present study demonstrates important and sexually dimorphic roles in disease-relevant tissues for murine *C2CD4B*, but not *C2CD4A* (Supp. Fig.17). These include contributions of the former gene to body weight control and insulin secretion (males) and to pituitary function and insulin secretion (females). Our data also highlight differences in the likely roles of these genes in humans *versus* other species.

## Author contributions

SNMG, GAR, NN and MV designed the research. SNMG, IC, PC, AMS, GP, SJM, EG, MH, BO, MV, NF, MD, ZM, XL, FLCD and ET performed experiments. AMS, DW, TJP, NN, PG, DAJ, AM and HK contributed new reagents/analytical tools. IL provided project licence. SNMG and GAR drafted and/or wrote the manuscript. GAR and Diabetes UK provided funding. GAR supervised the work.

We thank Dr. Ines Cebola, Imperial College London, for her comments and revision of the manuscript. We also thank Dr. Stephen Rothery and the Imperial College FILM facility for training and use of the wide-field and confocal microscope. We also thank Hermine Muniangi Muhitu for quantification of mouse islet β cell mass and Xiaomeng Li for plasmid cloning.

## Guarantor Statement

G.A.R. serves as the guarantor of this work

## Prior Presentation information

This work has been presented at Diabetes UK Annual Conference Manchester (2017), Diabetes UK APC (2018), EASD Annual Conference Berlin (2018) conferences.

## Abbreviations

FSH: follicle-stimulating hormone
GFP: green fluorescent protein
GWAS: genome-wide association study
IMPC: International Mouse Phenotyping Consortium
LH: luteinizing hormone
KREBH: Krebs-HEPES-bicarbonate buffer
RC: regular chow
HFD: high fat diet
T2D: Type 2 diabetes

