## Supplemental Methods for "Sexually dimorphic roles for the type 2 diabetes-associated *C2cd4b* gene in murine glucose homeostasis"

**Reagents**

Mouse monoclonal anti-FLAG antibody (Sigma-Aldrich, F1804-200UG), guinea-pig polyclonal anti-insulin (DAKO, ready to use, IR002), mouse monoclonal anti-glucagon antibody (Sigma-Aldrich, G2654-100UL), rabbit polyclonal antibody anti-TGN46 (Abcam, Ab16059), rat monoclonal anti-LAMP-1 antibody (Santa Cruz, Sc-19992) were used for immunostaining of fixed cells and/or paraffin sections. Alexa Fluor 488 goat anti-guinea-pig (A-11073), Alexa Fluor 532 goat anti-rabbit (A11009) and Alexa Fluor 488 goat anti-mouse (A11029) were used as secondary antibodies from Invitrogen.

***C2cd4a* and *C2cd4b* mouse lines**

Both strains were maintained on a C57BL/6N background. In the *C2cd4a* strain, 1742 bp were deleted from exon 2 and in *C2cd4b* strain, exon 2 was deleted. Heterozygous animals were used as breeding pairs. Genotyping was carried out according to the IMPC protocols.

Mice were housed in a pathogen-free facility with a 12-hour light/dark cycle and had free access to standard mouse chow diet and water. Animals fed with high-fat and -sucrose diet contained 58% fat and 25% carbohydrate, provided from Research Diets, Cat. No. D12331. Animals were maintained on and had free access to this diet from 6 weeks of age.

**Immunoprecipitation and mass spectrometry**

MIN6 cells grown in standard culture conditions (1) were transfected in duplicates with 1 µg FLAG-tagged C2CD4A, C2CD4B or with FLAG tag only expressing plasmids using Lipofectamine 2000. Immunoprecipitation and AP-MS analysis was performed as previously described (2) with minor modifications.

Samples were processed with on-bead digestion using 1M urea followed by Trypsin Gold (Promega), acidified with TFA and de-salted using reversed-phase spin tips (Glygen Corp). Dried peptides were subjected to LC-MS/MS analysis using an Ultimate 3000 nano HPLC coupled to a Q-Exactive mass spectrometer (Thermo Scientific) via an EASY-Spray source. Peptides were loaded onto a trap column (Acclaim PepMap 100 C18, 100µm × 2cm) at 8µL/min in 2% acetonitrile, 0.1% TFA. Peptides were then eluted on-line to an analytical column (Acclaim Pepmap RSLC C18, 75µm × 75cm). Data was processed using the MaxQuant (3) software platform (v1.6.2.3), with database searches carried out by the in-built Andromeda search engine against the Uniprot *Mus musculus* database (downloaded – 4th January 2018, entries: 83,123). A reverse decoy database approach was used at a 1% FDR for peptide spectrum matches and protein identifications. Search parameters included: maximum missed cleavages set to 2, variable modifications of methionine oxidation, protein N-terminal acetylation, asparagine deamidation and cyclisation of glutamine to pyroglutamate. Label-free quantification was enabled with an LFQ minimum ratio count of 2.

Proteins were filtered for those containing at least 2 razor and unique peptides, identified in all replicates with a minimum of a 2-fold increase in abundance compared with control (FLAG-tag only) immunoprecipitates and ranked by average intensity across replicates.

**Immunofluorescence and imaging for sub-cellular localisations**

Cells were cultured on coverslips and 12 or 24 h post transfection were fixed in 4% [wt/vol.] PFA (Sigma-Aldrich), and incubated with primary antibody (mouse anti-FLAG, Sigma-Aldrich, 1:1000). Samples were incubated with secondary antibody (Alexa Fluor 488 anti-mouse, Invitrogen, 1:1000) for 2 h at room temperature. A Nikon ECLIPSE Ti spinning disk microscope was used to collect images. A x60 oil immersion objective was used for localisation and co-localisation experiments. For co-localisation with ER experiments a confocal inverted Zeiss LSM-780 microscope was used to capture images.

**Zebrafish maintenance and generation of transgenic lines**

Zebrafish (*Danio rerio*) were raised and cared for according to standard protocols (4). All animal work has been conducted according to national guidelines and all animal experiments described herein were approved by the ethical committee of the University of Liège (protocol number 13-1557).

The transgene rs7163757C-cfos:eGFP was constructed by introducing a 1303 bp region carrying the GWAS rs7163757-C upstream of a c-fos minimal promoter driving EGFP (pGW_cfos-EGFP) (5,6) via LR recombination. Purified rs7163757C-cfos:eGFP transgene was injected into 1–2-cell zebrafish embryos and the resulting GFP expression pattern analysed at different time points during development (F0 analysis). The fluorescent injected fish were raised to adulthood and the offspring screened for fluorescence. GFP expression pattern of the transgenic line obtained was tested by whole mount immunohistochemistry as described in (7) using chicken anti-GFP (Aves Labs, 1:1000), guinea pig anti-Insulin (Dako,1:500) and fluorescently conjugated AlexaFluor antibodies (Invitrogen). Fluorescent images were acquired with a Leica SP5 confocal microscope.

**Pericardial glucose injection and live imaging in zebrafish larvae**

To record glucose-stimulated calcium dynamics in β cells *in vivo,* heterozygous animals carrying a 1kb deletion in the *C2cd4a* promoter were in-crossed. The progeny was sorted for the presence of the *Tg(ins:GCaMP6s,cryaa:RFP)*$^{tud202Tg}$ reporter, which was previously introduced in the background of the *C2cd4a* promoter deletion. Glucose injections and live imaging were performed blindly for the genotype of the larvae. After imaging, each larva was genotyped, and the genotype was assigned to the corresponding images.

Glucose injection and live imaging of zebrafish primary islets were performed as described previously (8). The embryos were treated with 0.003% (200μM) 1-phenyl-2-thiourea to inhibit pigmentation, starting from 1 dpf. At 4.5 dpf, the larvae were anaesthetized with 0.4 g/l tricaine (MS-222) and mounted in 1% low melting point agarose containing 0.4 g/l tricaine in 35 mm glass bottom petri dishes (MatTek Corporation). After the agarose solidified, embryo water containing 0.4 g/l tricaine was added onto the embryos.

Live imaging was performed on a Zeiss LSM 780 inverted confocal microscope with a 40x water lens (Zeiss C-Apochromat 40x/1.2 W autocorr M27). The GCaMP6s signal was acquired using the 488 laser line. Image acquisition was performed using a single focal plane, with a speed of 150 ms per frame (6.66Hz) for 400 frames (60 seconds), and an XY resolution of 512 x 512 pixels.

**Homogeneous Time Resolved Fluorescence (HTRF) Assay**

Insulin Ultra-Sensitive Kit (Cisbio, ref. 62IN2PEH) was used according to the manufacturer's instructions to measure released or total insulin levels. In order to assess the final dilutions for samples, a test with several dilutions was carried out before measuring all samples collected. Each sample was measured in duplicate and incubated with europium cryptate and XL665 antibodies overnight before measuring the Förster Resonance Energy Transfer (FRET) efficiency.

**Immunofluorescence on pancreatic slices**

Animals were dissected at 24 or 25 weeks of age. Pancreata were fixed in 4% [wt/vol.] paraformaldehyde (PFA, Sigma-Aldrich) diluted in phosphate-buffered saline (PBS). Samples sent to the histology facility at Imperial College London were embedded in paraffin and sectioned at 5 µm thickness. Each section was 150 µm apart from the previous section. Primary antibodies (guinea-pig anti-insulin ready to use (Dako): not diluted, mouse anti-glucagon (Sigma-Aldrich): 1:1000), diluted in PBS (containing: 0.25% BSA, 0.25% Triton X-100), were applied overnight at 4°C. Slides were incubated with secondary antibodies (Alexa Fluor 488 goat anti-guinea-pig, Invitrogen, 1:1000, Alexa Fluor 532 goat anti-rabbit, Invitrogen,1:1000, DAPI, Roche 1:5000) for 2 h at room temperature (RT). ProLong Dimond Antifade Mountant, Life Technologies was used for mounting. An inverted widefield microscope with LED illumination Zeiss Axio Observer microscope (Zeiss Axio Observer Z1), from the Imperial College FILM facility, was used to collect images.

**Intracellular free [$Ca^{2+}$] measurements**

Imaging was performed essentially as described (9). In brief, 24 h after islet isolation, 20 islets/acquisition were incubated for 45 min. in fluo2-AM (10 µM; Teflabs) diluted in a KREBH buffer solution containing 3 mM glucose. A Nipkow spinning disk head microscope was used to capture the fluorescent signals. Islets were maintained at 35°C to 36°C and continuously irrigated with KREBH buffer solution aerated with 95% $O_2$ and 5% $CO_2$. On average, 8-10 islets were imaged in each field of view. Two acquisitions were performed per animal. Images were analysed using ImageJ software (URL: https://imagej.nih.gov/ij/index.html) by measuring the fluorescence over time. The Pearson product moment correlation analysis was performed for every possible cell pair to assess inter-cellular connectivity (8).

**Sample preparation for RNA sequencing**

Isolated islets from 5 male mice/genotype were used for RNA purification. DNase treatment was performed using TURBO DNase (Invitrogen) according to the manufacturer's instructions. RNA quantity and integrity were assessed using an RNA 6000 Nano Kit (Agilent) and an Agilent 2100 Bioanalyzer. mRNA enrichment was achieved from 0.8-1 µg of total RNA using a NEBNext Poly(A) mRNA Magnetic Isolation Kit (NEB). Generation of double stranded cDNA and library construction were performed using NEBNext Ultra II Directional RNA Library Prep Kit for Illumina (NEB). NeBNext Multiplex Adapters (NEB) was used to perform ligation of the adapters. Each library was subsequently size selected with SPRIselect Beads (Beckman Coulter). The Adaptor ligated DNA was PCR amplified using NEBNext Ultra II Q5 Master Mix and Universal i5 and i7 primers provided in the NEBNext Kits.

Sequencing was performed by the Imperial BRC Genomics Facility as 75bp paired end reads on a HiSeq4000 according to Illumina specifications. FASTQ files were generated for each sample (5 WT and 4 null mice) and initial data quality checks of the raw sequence data were performed. Reads were then mapped to the mouse transcriptome (GRCm38, cDNA and ncRNA) using Salmon (10). Total RNA profiles were consistent and generated ~20-40 million reads mapping to Ensembl genes per sample. DESeq2 (v1.20.0) (11) with DESeq2-default normalization method and adjusted p-value threshold <0.1 was used for differential expression analysis in R using relevant BioConductor packages (12).

**Whole-cell voltage-clamp electrophysiology**

Wild type control and *C2cd4b* knockout mouse islets were dispersed into single cells by gently titration for 1 minute in 0.005% trypsin and cultured overnight in RPMI-1640 medium (RPMI) with 11 mM glucose supplemented with 15% fetal bovine serum (FBS), 100 IU·$ml^{-1}$ penicillin, and 100 mg·$ml^{-1}$ streptomycin at 37°C, 5% $CO_2$. Patch electrodes (3-4 MΩ) were backfilled with intracellular solution containing (mM) 102.0 CsCl, 10.0 TEA-Cl, 10.0 EGTA, 3.0 $Na_2$ATP, and 5.0 HEPES (pH 7.25 adjusted by CsOH). Voltage-clamp electrophysiology was performed on β-cells in extracellular buffer containing (mM) 119.0 NaCl, 4.7 KCl, 2.0 $CaCl_2$, 1.2 $MgSO_4$, 1.2 $KH_2PO_4$, 10.0 HEPES, and 17.0 glucose (pH 7.35 adjusted by NaOH). After forming a tight seal between the patch pipette and β-cell (seal resistance > 1 GΩ), whole-cell access was established and the bath solution was exchanged (3 minutes; 2 mL/minute flowrate) with extracellular buffer containing (mM) 82.0 NaCl, 5.0 CsCl, 30.0 $CaCl_2$, 1.0 $MgCl_2$, 0.1 EGTA, 20.0 TEA-Cl, 0.1 tolbutamide, and 17.0 glucose (pH 7.35 with NaOH). Starting from a holding potential of -80, VDCC currents were generated through application of sequential 10 mV depolarizing steps ranging from -70 to 70 mV (500 ms); membrane potential was held at -80 mV for 7.5 seconds between each voltage step. Linear leak currents were subtracted using a P/4 protocol. VDCC currents were normalized to cell capacitance and normalized peak VDCC currents plotted as a function of applied voltage.
