## Supplemental Tables for "Sexually dimorphic roles for the type 2 diabetes-associated *C2cd4b* gene in murine glucose homeostasis"

### Supplementary Tables

| Gene | Expression | Tissue | Publication | Expression | Tissue | Publication |
|---|---|---|---|---|---|---|
| Murine *C2cd4a* | 6.98 (RPKM) | β | Benner *et al.* (20) | 5.5 (RPKM) | Mouse pancreatic islets | Kone *et al.* (21) |
| Murine *C2cd4b* | 71.6 (RPKM) | | | 40.7 (RPKM) | | |
| Human *C2CD4A* | 57.9 (RPKM) | β | Benner *et al.* (20) | 18.5 (TPM) | β | Blodgett *et al.* (23) |
| Human *C2CD4B* | 52.8 (RPKM) | | | 15.7 (TPM) | | |

Supplementary Table 1. Expression of *C2CD4A* and *C2CD4B* in human and mouse islets and β-cells. While both *C2CD4A* and *C2CD4B* are expressed at the same level in human β-cells, in mouse *C2cd4b* is more predominantly expressed compared to *C2cd4a*. β=FACS purified β-cells.

| Gene name | Base Mean | log2FoldChange | Fold Change | p value | Adjusted p value |
|---|---|---|---|---|---|
| C2cd4b | 1139.51706 | -2.415152232 | 0.187485089 | 1.36E-44 | 3.81E-40 |
| C2cd4a | 118.027192 | 0.814689614 | 1.758919692 | 2.86E-06 | 0.040127223 |
| Cpsf1 | 1904.77052 | -0.578923095 | 0.669463314 | 7.06E-06 | 0.066105298 |
| C3 | 152.141699 | 0.741957418 | 1.672443436 | 3.45E-05 | 0.241943108 |
| Rpph1 | 140.612162 | -0.703996659 | 0.613869264 | 5.77E-05 | 0.323789867 |
| Gm45322 | 417.49085 | 0.620112657 | 1.536995197 | 0.00011 | 0.504901364 |
| mt-Nd4l | 3597.92578 | 0.680785322 | 1.603012107 | 0.000126 | 0.504901364 |
| Emid1 | 454.688167 | -0.607583874 | 0.656294899 | 0.000149 | 0.524402308 |
| Crb1 | 108.03916 | -0.647844838 | 0.638233023 | 0.000268 | 0.766730644 |
| Gm10076 | 52.4812133 | -0.547066149 | 0.684410525 | 0.000273 | 0.766730644 |

Supplementary Table 2. Effect of *C2cd4b* deletion on islet gene expression. The gene expression levels from isolated islets of four male mice per genotype were assessed by RNA-seq. Table presents gene expressions from high to low adjusted p values. The top ten genes with the highest adjusted p-values were selected in the above table. Highlighted in yellow are the genes with significant adjusted p-values. A significant reduction in *C2cd4b* expression levels was observed comparing islets from *C2cd4b* WT to null mice. *C2cd4a* expression was increased in *C2cd4b* null mice.

| Name | Unique peptides | Average intensity | FC over control | Function |
|---|---|---|---|---|
| C2cd4b;C2cd4a | 4 | 1692900000 | infinity | |
| 60S ribosomal protein L36 (Rpl36) | 7 | 408547500 | 2.31651774 | Ribosomal protein |
| Bone morphogenetic protein 1 (Bmp1) | 14 | 323217500 | 4.528805647 | Formation of cartilage in vivo |
| Clusterin beta/alpha chain (Clu) | 6 | 242922500 | infinity | Basic biological events such as cell death, tumour progression, and neurodegenerative disorders |
| Sequestosome-1 (Sqstm1) | 4 | 238032500 | 4.041972241 | Scaffolding/adaptor protein in concert with TNF receptor-associated factor 6 to mediate activation of NF-kB |
| Glucagon;Glicentin (Gcg) | 2 | 149391750 | infinity | Glucagon, glucose homeostasis |
| Ttc13 | 9 | 142825000 | 2.147951497 | A protein coding gene, unknown function |
| Procollagen-lysine,2-oxoglutarate 5-dioxygenase 3 (Plod3) | 8 | 137387500 | infinity | Catalyses the hydroxylation of lysyl residues in collagen-like peptides |
| Hyaluronan and proteoglycan link protein 4 (Hapln4) | 4 | 129820000 | infinity | Extracellular matrix structural constituent and hyaluronic acid binding |
| Torsin-2A;Prosalusin;Salusin-beta (Tor2a) | 3 | 107971000 | infinity | Increases intracellular [$Ca^{2+}$], induces cell mitogenesis |
| U5 small nuclear ribonucleoprotein (Snrnp40) | 4 | 103415000 | infinity | Component of the U5 small nuclear ribonucleoprotein (snRNP) particle |
| MAP7 domain-containing protein 1 (Map7d1) | 3 | 102242500 | infinity | Structural molecule activity |
| Nucleolar protein 14 (Nop14) | 4 | 91338750 | infinity | Pre-18s rRNA processing and small ribosomal subunit assembly |
| DNA topoisomerase 2-beta (Top2b) | 4 | 88047000 | infinity | Chromosome condensation, chromatid separation, and the relief of torsional stress |
| Proprotein convertase subtilisin/kexin type 9 (Pcsk9) | 6 | 86949000 | infinity | Processes protein and peptide precursors trafficking |
| Receptor-type tyrosine-protein phosphatase N2 (Ptprn2) | 4 | 82972000 | infinity | Required for normal accumulation of secretory vesicles in hippocampus, pituitary and pancreatic islets |

Supplementary Table 3. C2CD4A interacting proteins detected by mass-spectrometry. Top 15 predicted interacting proteins for C2CD4A in MIN6 cells. The given function(s) of the identified protein was adopted from GeneCards.org.

| Name | Unique peptides | Average intensity | FC over control | Function |
|---|---|---|---|---|
| Neuroendocrine convertase 2 (Pcsk2) | 18 | 2188050000 | 2.365537387 | Is a protease that processes protein and peptide precursors trafficking through the secretory pathway |
| Renin receptor (Atp6ap2) | 8 | 1197847500 | 4.624034467 | Associated with the transmembrane sector of the V-type ATPases |
| Carboxypeptidase E (Cpe) | 12 | 852947500 | 2.488765327 | Involved in the biosynthesis of peptide hormones and neurotransmitters, including insulin |
| C2cd4a; C2cd4b | 4 | 852000000 | infinity | |
| Clusterin beta chain;Clusterin alpha chain (Clu) | 6 | 364380000 | infinity | Basic biological events such as cell death, tumour progression, and neurodegenerative disorders |
| N-acetylglucosamine-1-phosphotransferase subunit gamma (Tce7;Gnptg) | 8 | 361852500 | 3.757036085 | Sub-unit of an enzyme that catalyses the formation of mannose 6-phosphate (M6P) in the Golgi apparatus |
| Importin-5 (Ipo5) | 5 | 274114500 | infinity | Nucleoplasmic transport of proteins containing NLS |
| Carbohydrate sulfotransferase 11 (Chst11) | 4 | 238460000 | 2.402356026 | Catalyses the transfer of sulphate to position 4 of the N-acetylgalactosamine (GalNAc) residue of chondroitin |
| Glucagon (Gcg) | 2 | 217025000 | infinity | Glucagon, glucose homeostasis |
| Sequestosome-1 (Sqstm1) | 4 | 211023500 | 3.868217344 | Regulates activation of the nuclear factor kappa-B (NF-kB) signalling pathway |
| Bone morphogenetic protein 1 (Bmp1) | 14 | 200115000 | 3.837129646 | Formation of cartilage in vivo |
| Procollagen-lysine,2-oxoglutarate 5-dioxygenase 3 (Plod3) | 8 | 195605000 | infinity | Autophagy receptor required for selective macro-autophagy (aggrephagy). |
| Hyaluronan and proteoglycan link protein 4 (Hapln4) | 4 | 144687500 | infinity | A protein involved in Integrin Pathway and ERK Signalling |
| Receptor-type tyrosine-protein phosphatase N2 (Ptprn2) | 4 | 107998500 | infinity | Required for normal accumulation of secretory vesicles in hippocampus, pituitary and pancreatic islets (39–41) |

Supplementary Table 4. C2CD4B interacting proteins detected by mass-spectrometry. Top 15 predicted interacting proteins for C2CD4B in MIN6 cells. The given function(s) of the identified protein was adopted from GeneCards.org.
