## Supplemental figures for "Sexually dimorphic roles for the type 2 diabetes-associated *C2cd4b* gene in murine glucose homeostasis"

```
                                             C2+ binding site one

Topology          Strand1       Strand2            Strand3          Strand4
Consensus aa                    L V    A  L        DPYV   L          K
 PKCα          KRGRIYLKAEVADEKLHVTVRDAKNLIPMDPNGLSDPYVKLKLIPDPK-NESKQKTKTIR    59
 C2CD4A        ---------------LRLRLLRAESPAGG-APGPRAVSCRLSLVLRPPGTALRQCSTVVG    44
 C2CD4B        ---------------LRLRLLRAESLFGG-APGPRAVRCRLSLVLRPPGTARWQCSAVVG    44
                              *:: :  *:.      *      :*.*:  *   .   *  :  .:

                                             C2+ binding site two

Topology         Strand5             Strand6           Strand7          Strand8...
Consensus aa      NP WNE F F               L   V D D        G                W
 PKCα          STLNPQWNESFTFKLKPSDKDRRLSVEIWDWDRTTRNDFMGSLSFGVSELMKMPASGWYK    119
 C2CD4A        RSRKASFDQDFCFDGLSEDEVRRLAVRVKARDEGRGRERGRLL--GQGE-----------    91
 C2CD4B        RSRKASFDQDFCFDGLSEDEVRRLAVRVKARDEGRGRDRGRLL--GQGE-----------    91
                : : .:::.* *.   .*: ***:*.:   *.   .:    *  * .*

Topology       --
Consensus aa   L

 PKCα          LLNQE          124
 C2CD4A        -----          91
 C2CD4B        -----          91
```

Supplementary Figure 1. Amino acid sequence alignment of the C2 domains of hPKCα, hC2CD4A and hC2CD4B. The secondary structure is schematically indicated above the sequence as hashed lines, which indicate β-strands in synaptotagmin, a $Ca^{2+}$ sensor protein. The consensus sequence present in >50% of the C2 domains from 65 previously published C2 domains are indicated in bold on top (PKC structure and topology adopted from Nalefski *et al,* 1996). Amino acids shown in bold and black are non-polar or aromatic while if shown in orange they are polar or charged amino acids in consensus C2 domains. Highlighted in grey are side chains which are predicted to coordinate $Ca^{2+}$ in synaptotagmin and PKCα. The amino acids shown in blue are conserved in PKCα; shown in green are identical aa sequences between C2CD4A and C2CD4B. (Figure generated using sequence alignment on NCBI).

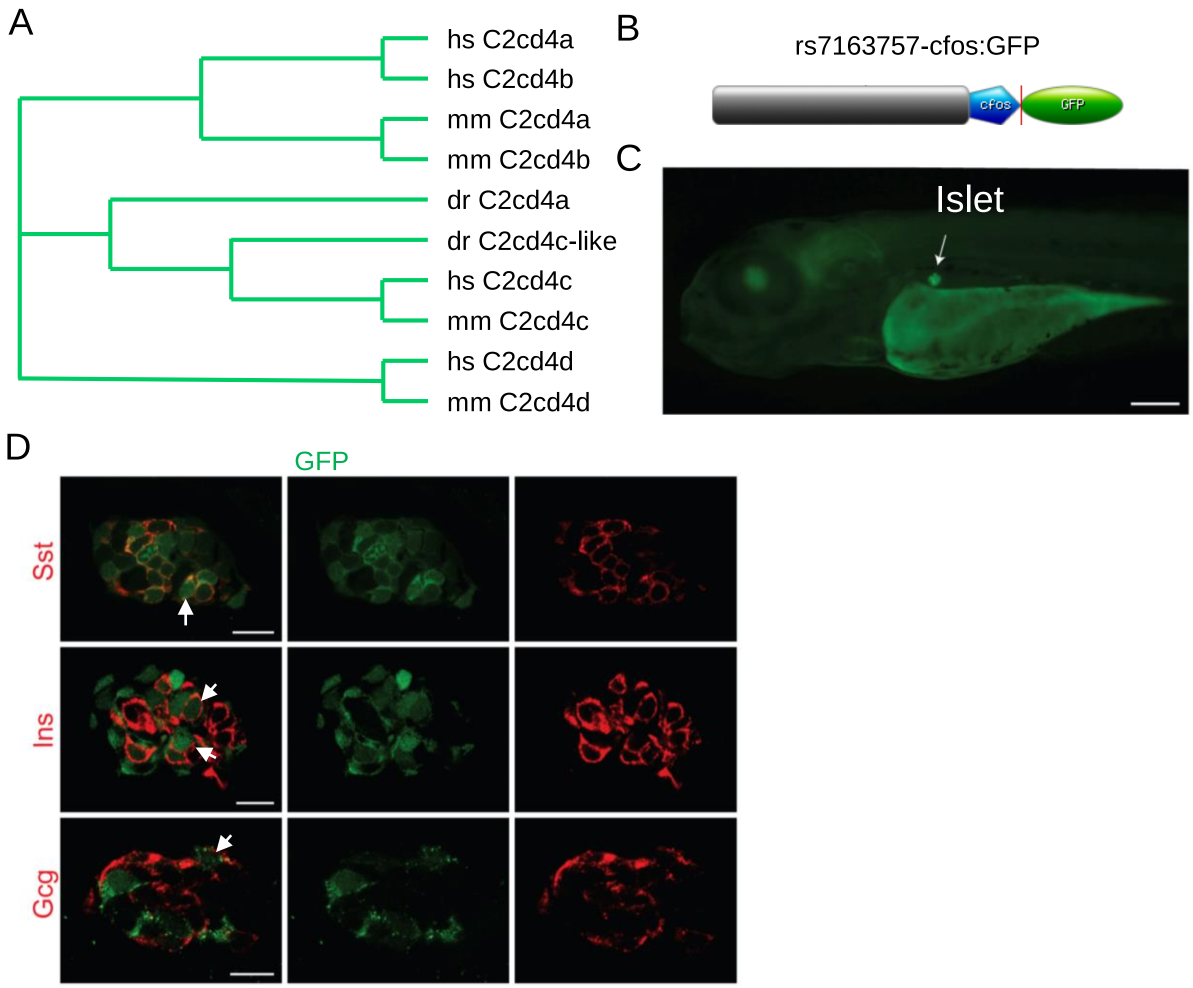


Supplementary Figure 2. The C2CD4 family and control of expression in zebrafish. A. The Phylogenetic tree was reproduced based on AlignX program of Vector NTI, using the full-length amino acid sequences of vertebrate members of the C2CD4 family. The numbers in parenthesis are their calculated evolutionary distances. dr-C2cd4a (ENSDARG00000061416, chr 25, previously called dr-C2cd4ab), dr-C2cd4c-like (ENSDARG00000079876.3, CU855878, chr 22). B. The rs7163757 region is able to direct the expression of GFP specifically in the endocrine pancreas and neuons in zebrafish. Schematic representation of the reporter construct rs7163757-cfos:GFP where GFP is under the control of a minimal promoter c-fos and of 1303 bp of the enhancer region containing the rs7163757 SNP. C. Binocular image of 4 dpf F2 transgenic line, rs7163757-cfos:GFP, revealing the expression of the GFP in the pancreas. Scale Bar : 200 µm. D. Whole-mount immunohistochemistry of 4 dpf rs7163757-cfos:GFP fish showing that GFP is expressed in the vast majority of somatostatin (sst)-positive cells, in most insulin-positive cells and in a few glucagon (gcg)-positive cells. Arrows indicate colocalization of dr-C2CD4A with the red channel. Scale Bar : 10 µm. Hs: Homo Sapiens, mm: mus musculus, dr: danio rerio.

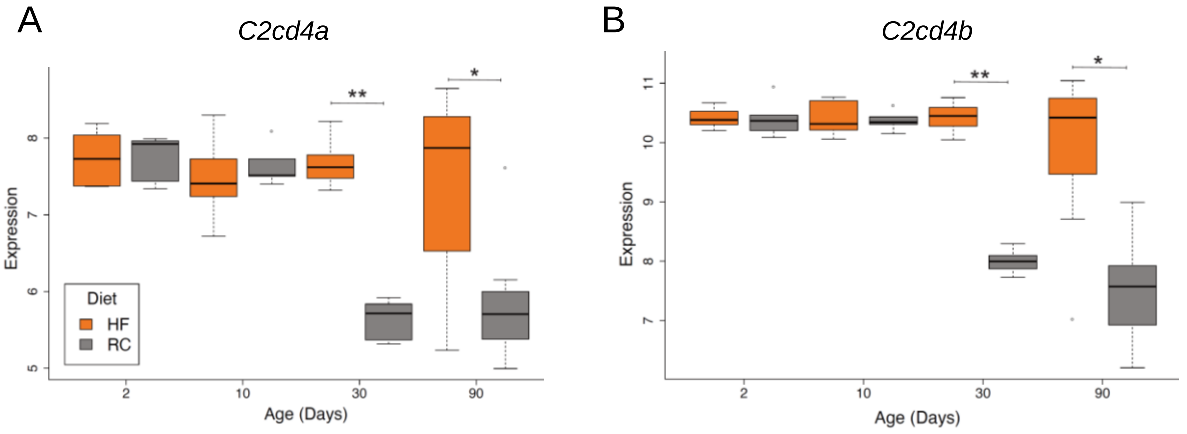


Supplementary Figure 3. Boxplots of *C2cd4a* and *C2cd4b* normalised expression level, see (31) for details, in DBA/2J mice fed a high fat (HF) or regular chow (RC) diet for 2, 10, 30 and 90 days. A. *C2cd4a* expression is significantly elevated in HF vs RC at 30 days (limma moderated t-test p-value=1.68e-10; Benjamini Hochberg adjusted p-value = 8.48e-9) and at 90 days (limma moderated t-test p-value=0.009; Benjamini Hochberg adjusted p-value = 0.029). B. *C2cd4b* expression is significantly elevated in HF vs RC at 30 days (limma moderated t-test p-value=4.32e-15; Benjamini Hochberg adjusted p-value = 4.58e-12) and at 90 days (limma moderated t-test p-value=0.0003; Benjamini Hochberg adjusted p-value= 0.0196). **adjusted p-value ≤0.01, *adjusted p-value ≤0.05.

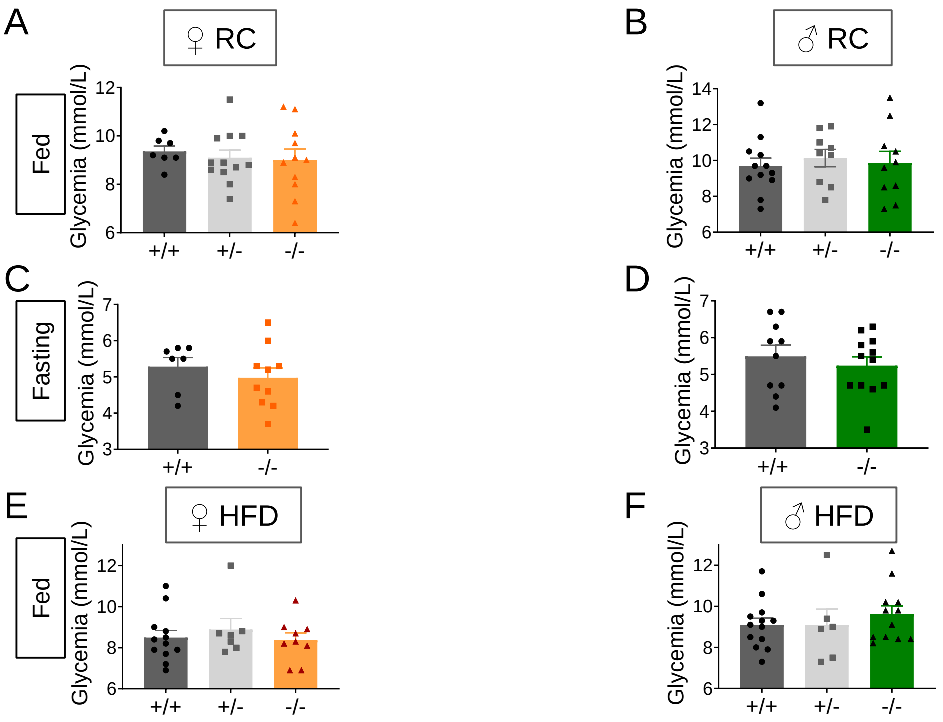


Supplementary Figure 4. *C2cd4b* mice glycemia. A-D. Fed and fasting glycemia in *C2cd4b*$^{+/+}$ and *C2cd4b*$^{-/-}$ mice on RC, at 18 and 20 weeks of age respectively (number of animals used for fed glycemia, $F^{+/+}$ n=7, $F^{+/-}$ n=12, $F^{-/-}$ n=11; $M^{+/+}$ n=12, $M^{+/-}$ n=9, $M^{-/-}$ n=10; fasting glycemia: $F^{+/+}$ n=7, $F^{-/-}$ n=10, $M^{+/+}$ n=10, $F^{-/-}$ n=12). E-F. Fasting glycemia in animals on HFD were measured at 23 weeks of age (number of animals used: $F^{+/+}$ n=9, $F^{-/-}$ n=5; $M^{+/+}$ n=5, $M^{-/-}$ n=7). Data were assessed for significance using an unpaired Student's t-test, or 2-way ANOVA where three genotypes were compared. Values represent means ± SEM.

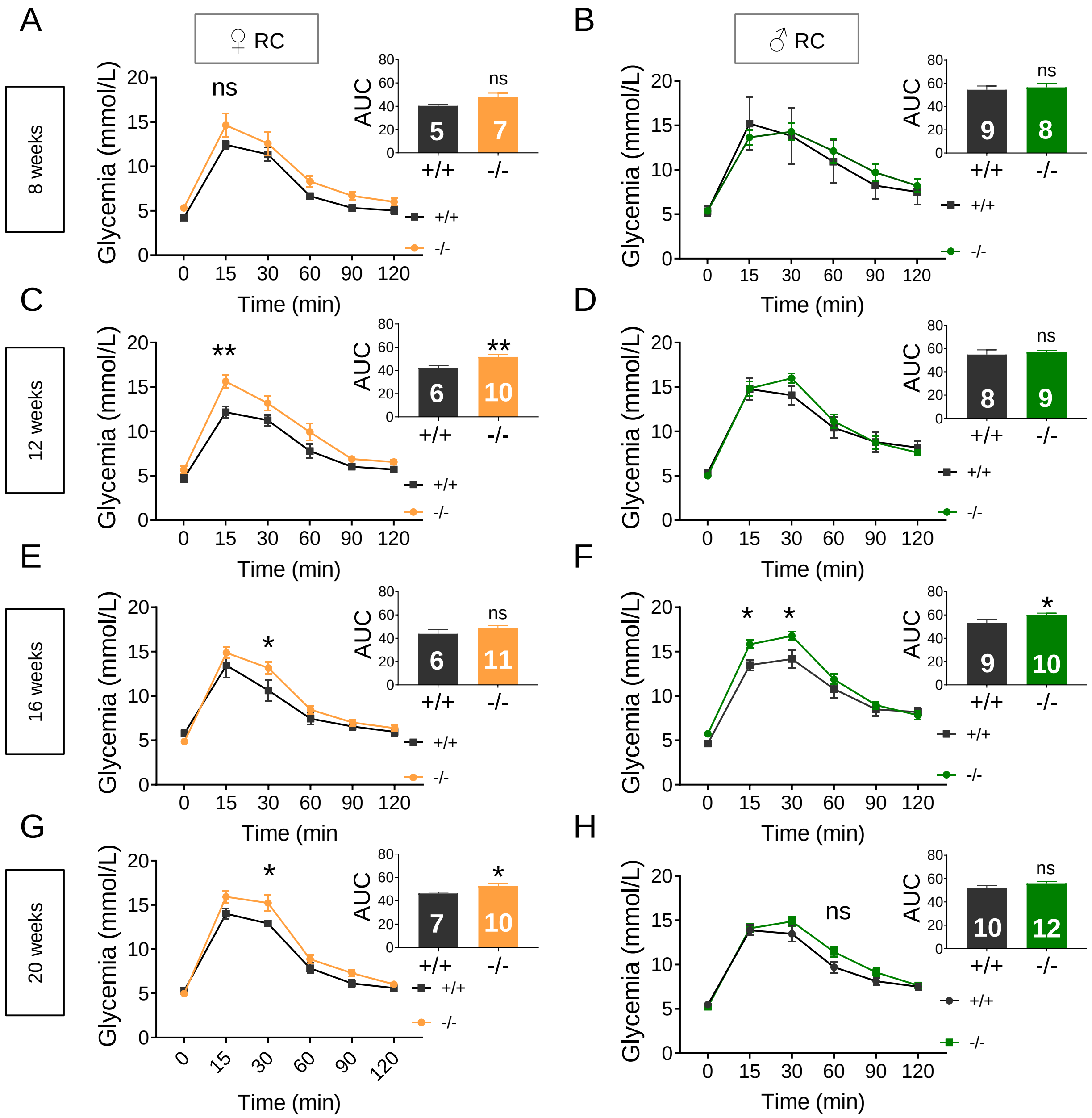


Supplementary Figure 5. Effect of deletion of *C2cd4b* on IPGGTs in mice maintained on RC. IPGTTs were performed on *C2cd4b* null male (B,D,F,H) and female (A,C,E,G) animals on RC at 8, 12, 16 and 20 weeks of age. *p<0.05, **p<0.01, ***p<0.001, data were assessed for significance using a 2-way ANOVA with Bonferroni's multiple comparison test. Inset: area under the curve (AUC) analysis; assessed for significance using an unpaired Student's t-test. Numbers in bar graphs represent the number of animals used (same number of samples used for glycemia and AUC graphs). Values represent means ± SEM.

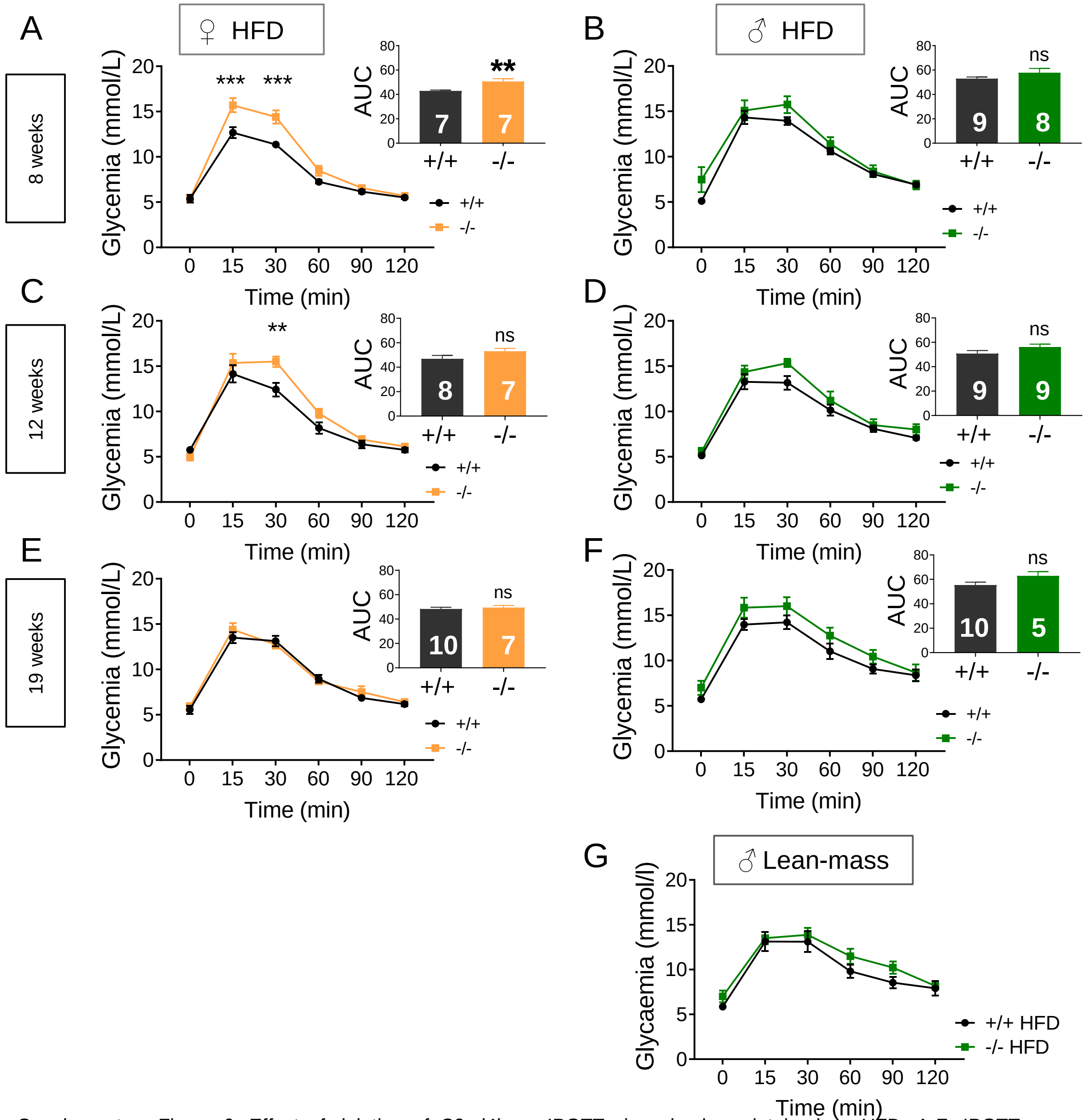


Supplementary Figure 6. Effect of deletion of *C2cd4b* on IPGTTs in animals maintained on HFD. A-F. IPGTTs were performed on *C2cd4b* null female (A,C,E) and male (B,D,F) animals on HFD at 8, 12, and 19 weeks of age. G. IPGTTs according to lean mass from *C2cd4b* male mice at 25 weeks of age. *p<0.05, **p<0.01, ***p<0.001, 2-way ANOVA with Bonferroni's multiple comparison test. Inset: area under the curve (AUC) analysis; data were assessed for significance using an unpaired Student's t-test. Numbers in bar graphs represent the number of animals used (same number of samples used for Glycemia and AUC graphs). Values represent means ± SEM.

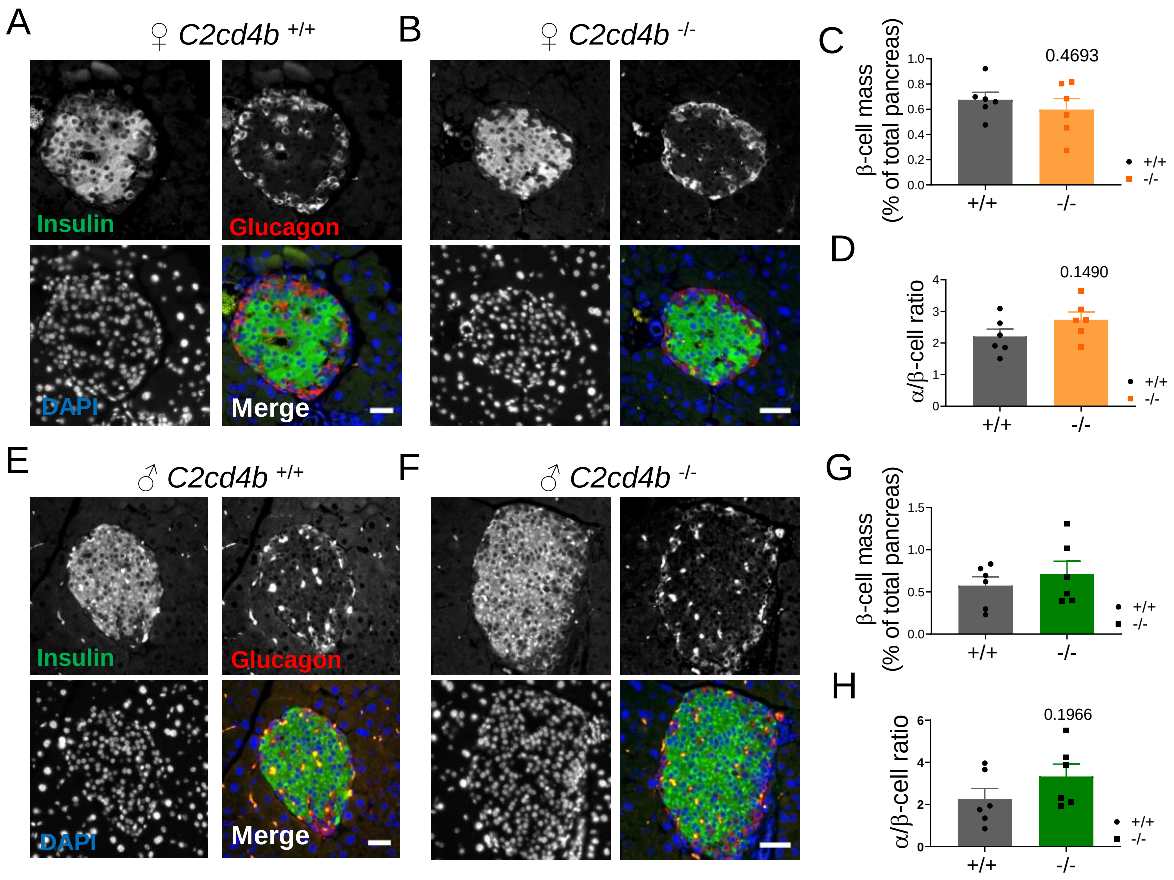


Supplementary Figure 7. β-cell mass in *C2cd4b* null mice. Data were collected from dissected pancreas from six mice per genotype at 24 weeks of age. For quantifications, three and two slides per female and male animals, respectively, were used. A,B,E,F. Immunohistochemistry was performed on slide sections with anti-insulin (shown in green), anti-glucagon (shown in red) antibodies, and DAPI (shown in blue). Scale bars= 30 µm. C,D,G,H. Area stained with insulin or glucagon antibodies were quantified to represent β-cell and α-cell mass, respectively, using ImageJ software. Data were assessed for significance using an unpaired Student's t-test. Values represent mean ± SEM.

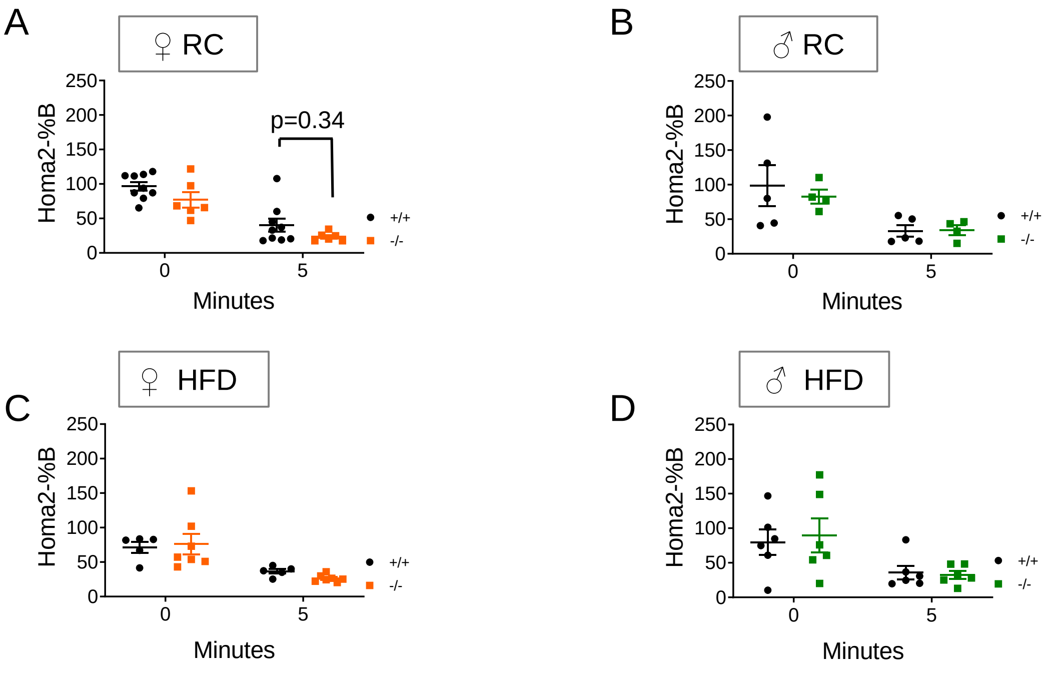


Supplementary Figure 8. Homeostatic model assessment analysis (HOMA2-%B) in *C2cd4b* mice. HOMA2-%B test was performed based on plasma insulin levels and glycemia at 0 and 5 min after injection with 3 g/kg body weight of glucose. Homeostatic model assessment analysis (HOMA2)-%B was measured using the HOMA Calculator from (https://www.dtu.ox.ac.uk/homacalculator/download.php). was used to calculate HOMA2-%B. A-B. HOMA2-%B was calculated using glycemia and plasma insulin measurements from *C2cd4b* female (A) and male (B) mice maintained on RC at 23 weeks of age. C-D. HOMA2-%B was calculated using glycemia and plasma insulin measurements from *C2cd4b* female (C) and male (D) mice maintained on RC at 19 weeks of age. *p<0.05, **p<0.01, ***p<0.001, 2-way ANOVA with Bonferroni's multiple comparison test. Data were assessed for significance using an unpaired Student's t-test. Values represent means ± SEM.

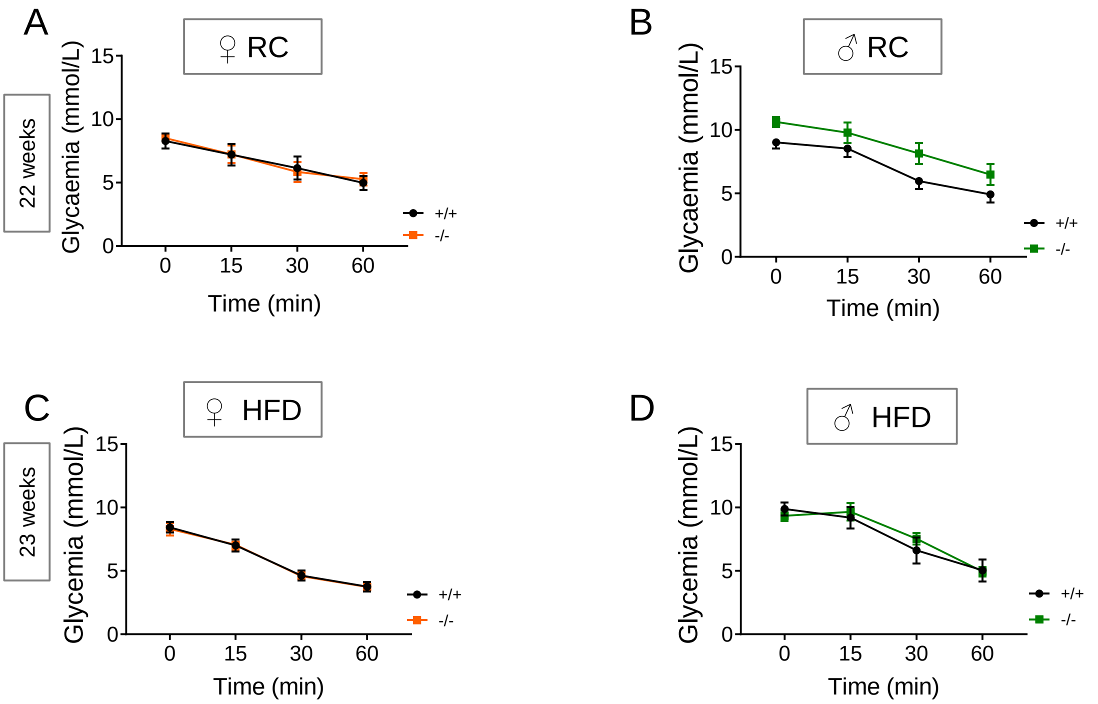


Supplementary Figure 9. Effect of deletion of *C2cd4b* on insulin sensitivity. *C2cd4b* null and WT male ($M^{+/+}$ n=9, $M^{-/-}$ n=13) and female ($F^{+/+}$ n=6, $F^{-/-}$ n=9) mice on RC were injected with a 1 or 0.75 Unit/kg body weight dose of insulin, respectively. On HFD, *C2cd4b* null and WT male (n=8/genotype) and female ($F^{+/+}$ n=11, $F^{-/-}$ n=7) mice were injected with 1.5 or 0.75 Unit/kg body weight dose of insulin. Glycemia was monitored over a one-h period. Deletion of *C2cd4b* had no effect on insulin sensitivity in either sex, examined under RC or HFD. Data were assessed for significance using a 2-way ANOVA with Bonferroni's multiple comparison test. Values represent mean ± SEM.

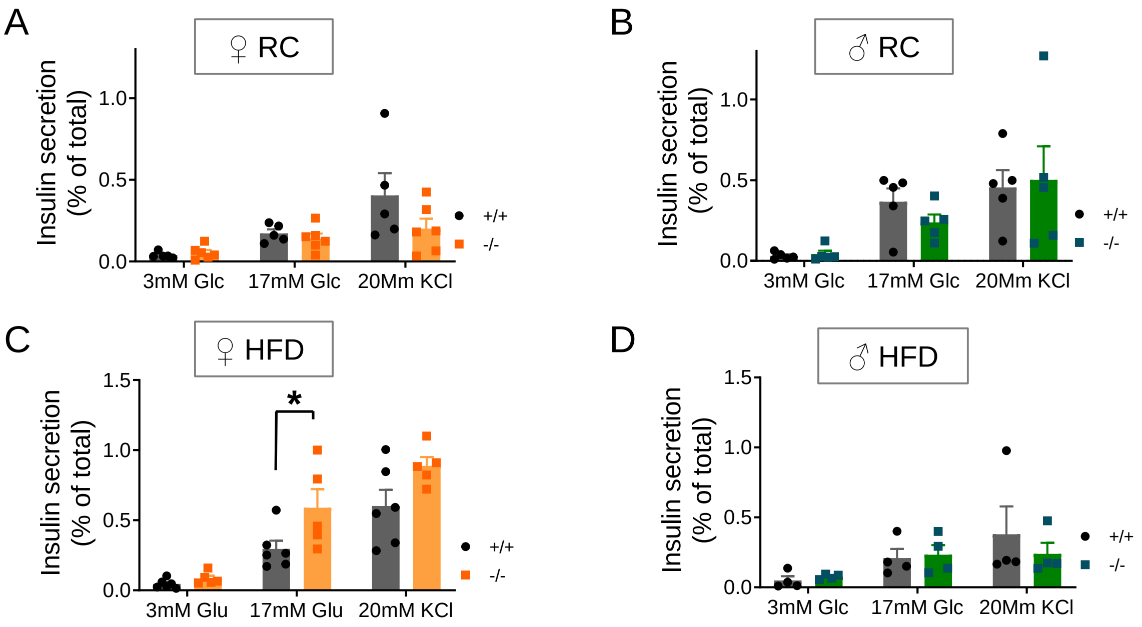


Supplementary Figure 10. Effect of deletion of *C2cd4b* on glucose or KCl-stimulated insulin secretion from isolated islets. Insulin secretion was measured in islets from *C2cd4b* mice maintained on RC (A-B) or HFD (C-D) at 24 weeks of age. (RC: $F^{+/+}$ n=5, $F^{-/-}$ n=6, M n=5/genotype; HFD: $F^{+/+}$ n=6 $F^{-/-}$ n=5, M n=4/genotype). * $p<0.05$, ** $p<0.01$, *** $p<0.001$, 2-way ANOVA with Bonferroni's multiple comparison test. Values represent mean ± SEM.

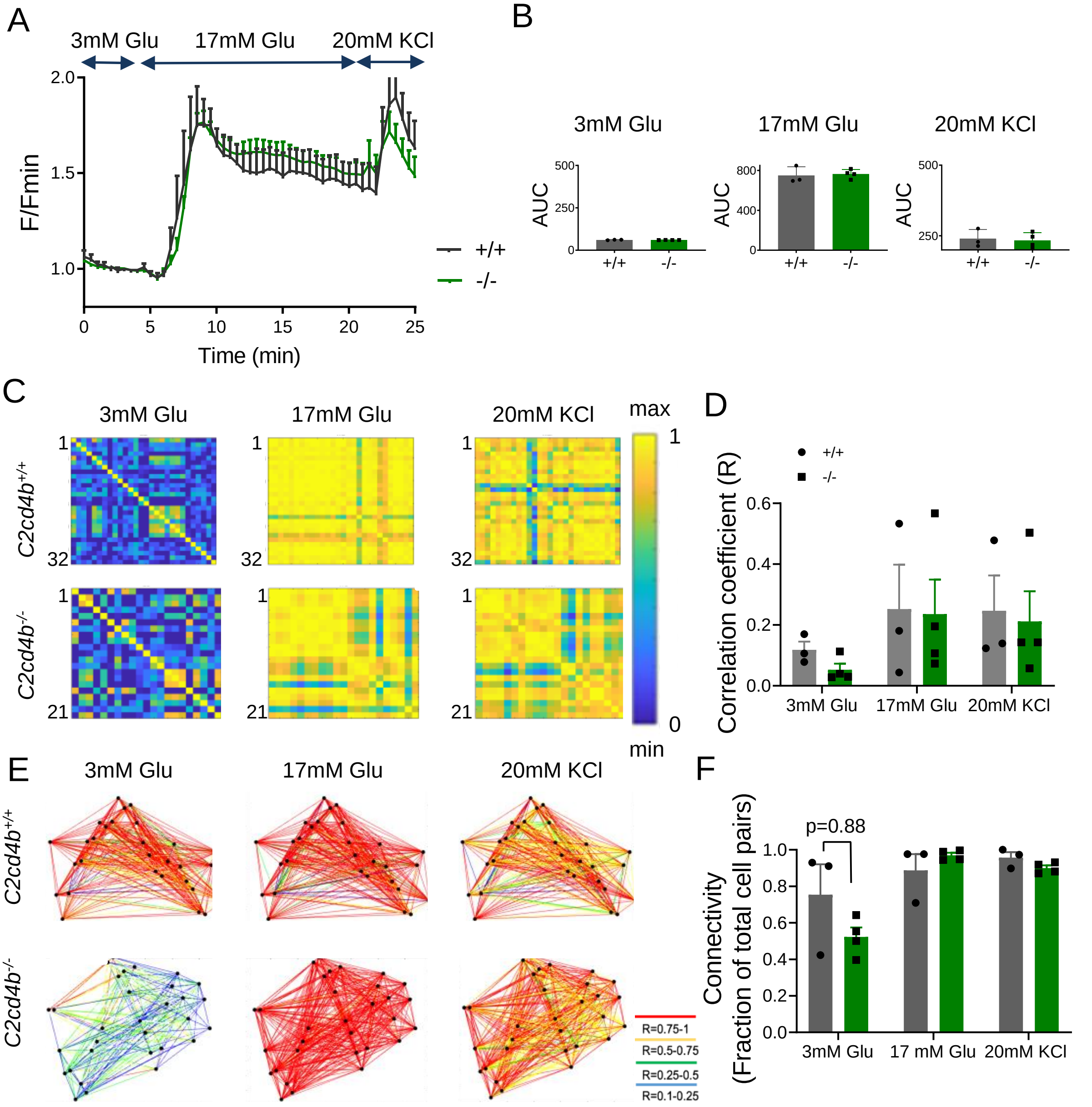


Supplementary Figure 11. Effect of deletion of *C2cd4b* on $Ca^{2+}$ dynamics in isolated islets. Intracellular $Ca^{2+}$ dynamics were assessed in isolated islets from females at 24 weeks of age maintained on RC, using spinning disk microscopy. A-B. No changes were observed in 17 mM glucose (Glu)- or 20 mM KCl-stimulated $Ca^{2+}$ dynamics between *C2cd4b* null and WT ($F^{+/+}$ n=3 and $F^{-/-}$ n=4 per experiments, two acquisitions were performed in each experiment and between 8-12 islets were assessed in each acquisition; assessed for significance using an unpaired Student's t-test). C-F. No significant changes in β-cell connectivity and in the correlation coefficient was observed between the null and WT islets (three islets assessed in duplicate for each genotype; data were assessed for significance using a 2-way ANOVA with Bonferroni's multiple comparison test. Values represent mean ± SEM.

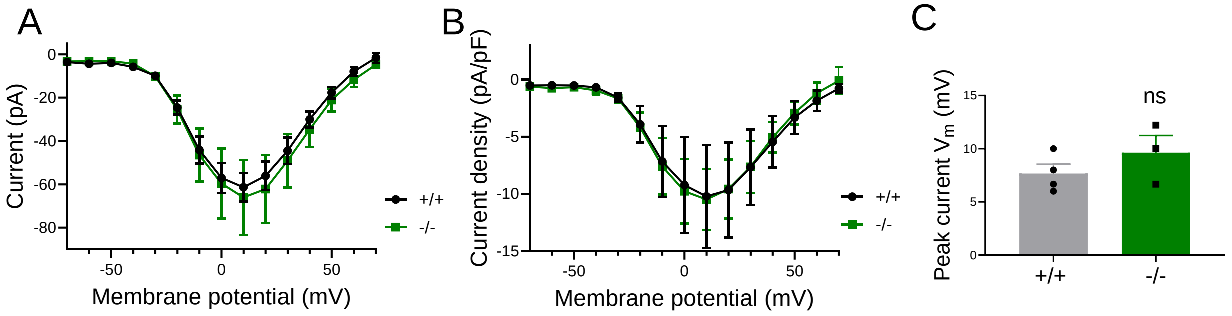


Supplementary Figure 12. *C2cd4b* deletion does not affect β cell voltage-dependent calcium channel activity. A. Average VDCC currents recorded from control *C2cd4b*$^{+/+}$ and *C2cd4b*$^{-/-}$ β cells in response to 10 mV steps from −70 to 70 mV (*C2cd4b*$^{+/+}$, n=18 β-cells (from 4 mice); *C2cd4b*$^{-/-}$ , n=26 cells (from 3 mice). B. Average WT and *C2cd4b* null β cell VDCC currents normalized to cell capacitance. C. average peak WT (gray) and *C2cd4b* null (green) β cell VDCC currents normalised to cell capacitance. Data were assessed for significance using an unpaired Student's t-test. Values represent mean ± SEM.

.

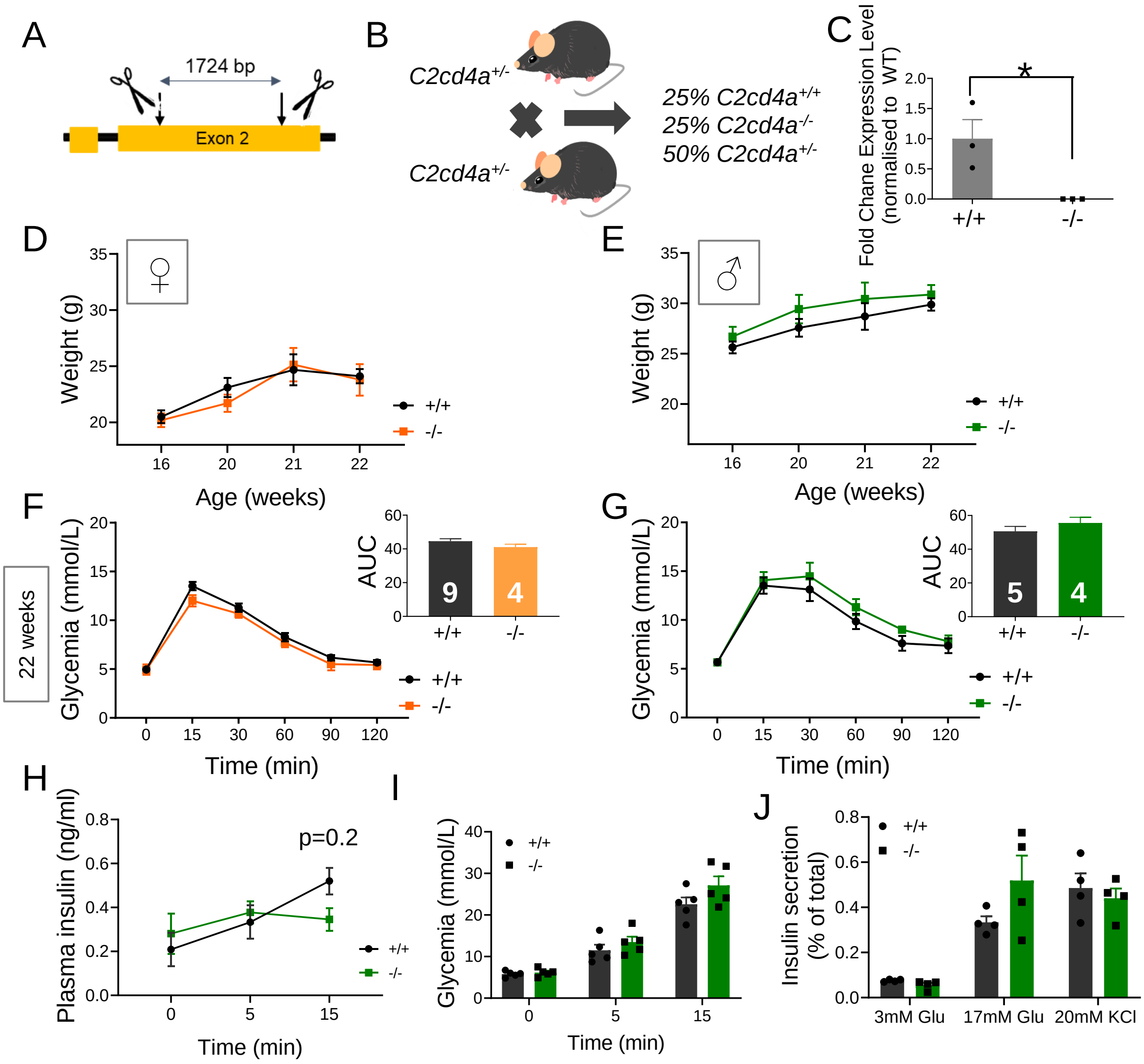


Supplementary Figure 13. *C2cd4a* null mice. A. *C2cd4a* mouse strain was generated by the IMPC using CRISPR/Cas9 to target 1724 bps from exon 2 of *C2cd4a* gene (See Methods). B. Inter-crossing of heterozygous mice resulted in the generation of wild type (WT, *C2cd4a*$^{+/+}$), heterozygous (*C2cd4a*$^{+/-}$) and null (*C2cd4a*$^{-/-}$) littermates. C. RT-q-PCR on RNA in isolated islets revealed a significant decrease in *C2cd4a* mRNA levels in homozygous animals (p=0.034), (n=3/genotype). *p<0.05, data were assessed for significance using an unpaired Student's t-test. D,E. Body weight of *C2cd4a* females (D) and males (E) at 16, 20. 21 and 22 weeks of age ($F^{+/+}$ n=4-11, $F^{-/-}$ n=4-9, $M^{+/+}$ n=2-8, $M^{-/-}$ n=3-6 animals per stage). Mixed-effect analysis, Tukey's multiple comparison test. F,G. IPGTTs were performed on *C2cd4b* female (F) and male (G) mice maintained on RC at 22 weeks of age. Inset: area under the curve (AUC) analysis, assessed for significance using an unpaired Student's t-test (same number of samples used for glycemia and AUC graphs). H,I. Intraperitoneal glucose tolerance tests were performed on *C2cd4a* null male animals maintained on RC at 21 weeks of age. Blood samples were collected, and glycemia measured at 5 and 15 min. after injection of glucose at 3 g/kg body weight. n=5/genotype. J. *In vitro* insulin levels in *C2cd4a* null *male* and WT mice maintained on RC. Measurements of insulin secretion were performed after stimulating isolated islets with 17 mM glucose or 20 mM KCl (n= 4/genotype). . *p<0.05, **p<0.01, ***p<0.001, data were assessed for significance using a 2-way ANOVA with Bonferroni's multiple comparison test. Values represent mean ± SEM.

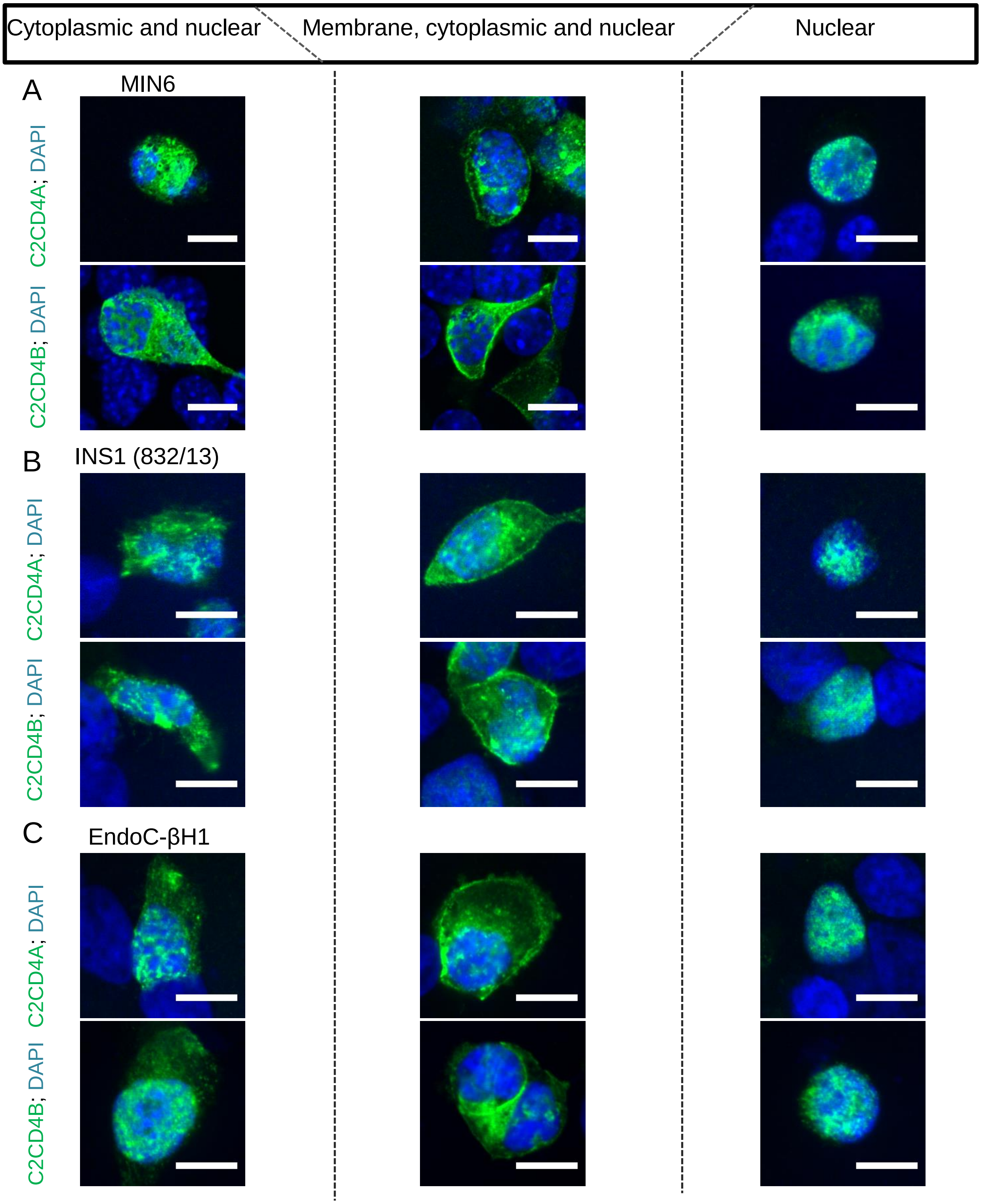


Supplementary Figure 14. Sub-cellular localisation of C2CD4A and C2CD4B in β-cells. Cells were transfected with either C2CD4A-FLAG or C2CD4B-FLAG tagged constructs and immunohistochemistry was performed using anti-FLAG (shown in green) antibody. Three main localisation patterns were observed by visualising C2CD4A (green) and C2CD4B (green) proteins in (A) MIN6, (B) INS1 (832/13) and (C) EndoC βH1 cells. DAPI is shown in blue. Scale bars=10 μm.

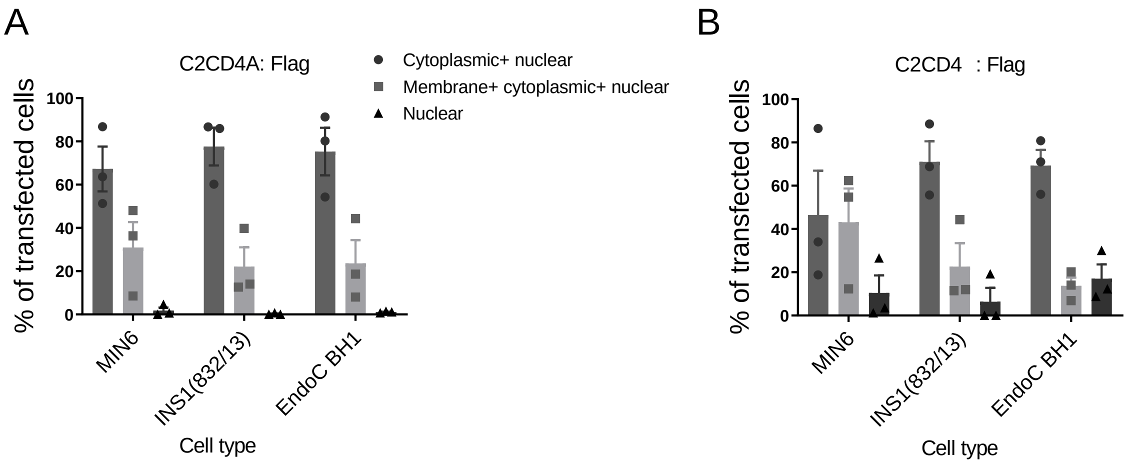


Supplementary Figure 15. Quantification of different localisation patterns in different β-cell types using C2CD4A: FLAG (A) and C2CD4B: FLAG-tagged (B) constructs. The quantifications were performed in three independent experiments. In each case, two separate cultures were analysed. Approximately 150 cells from two separate cultures were analysed in each experiment. No changes were detected in the proportions of the different localisation patterns between different cell types. Data were assessed for significance using a 2-way ANOVA with Bonferroni's multiple comparison test. Values represent means ± SEM.

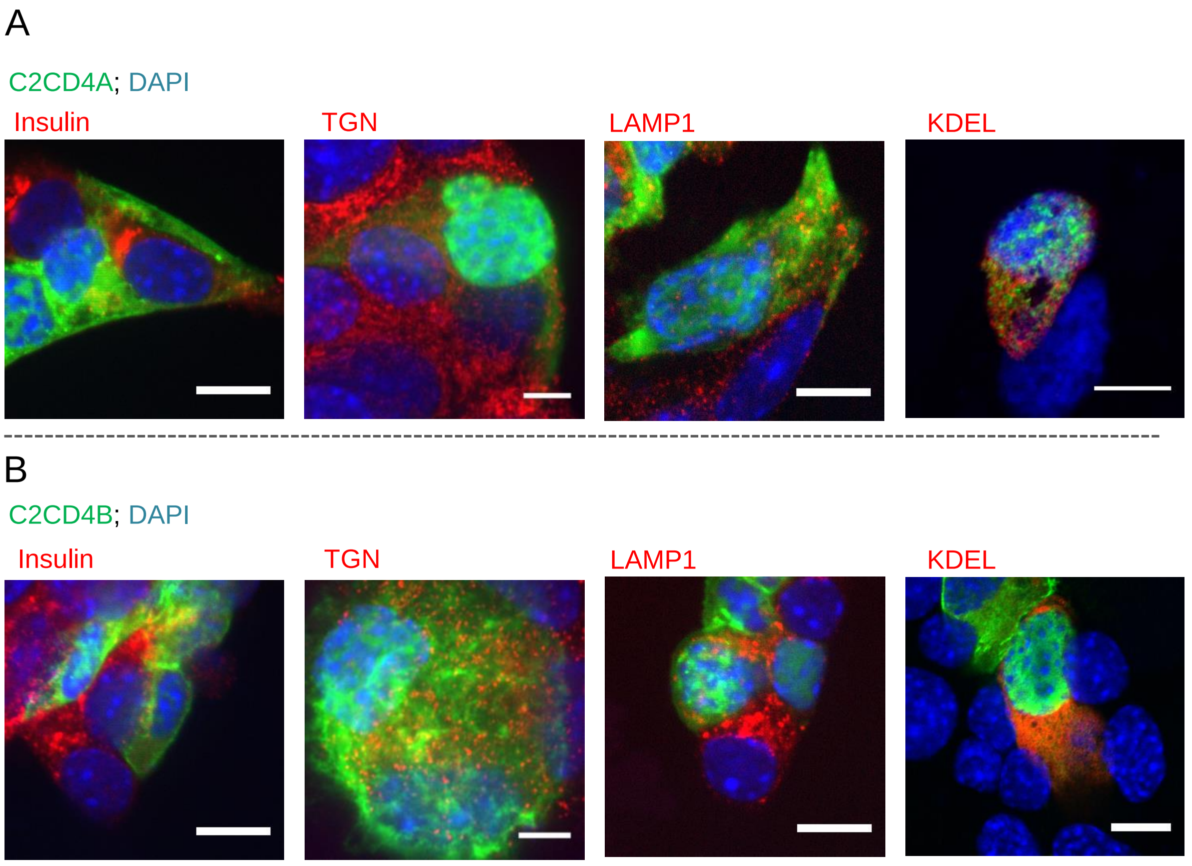


Supplementary Figure 16. Sub-cellular co-localisation of C2CD4A and C2CD4B in MIN6 cells. A. Sub-cellular co-localisation of C2CD4A (in green) with insulin, TGN (TGN46), lysosomes (LAMP1) and ER (KDEL) (all in red). B. Sub-cellular co-localisation of C2CD4B (in green) with insulin, TGN, lysosomes (LAMP1) and ER (KDEL) (all in red). DAPI staining shown in blue. Scale bars= 10 µm.

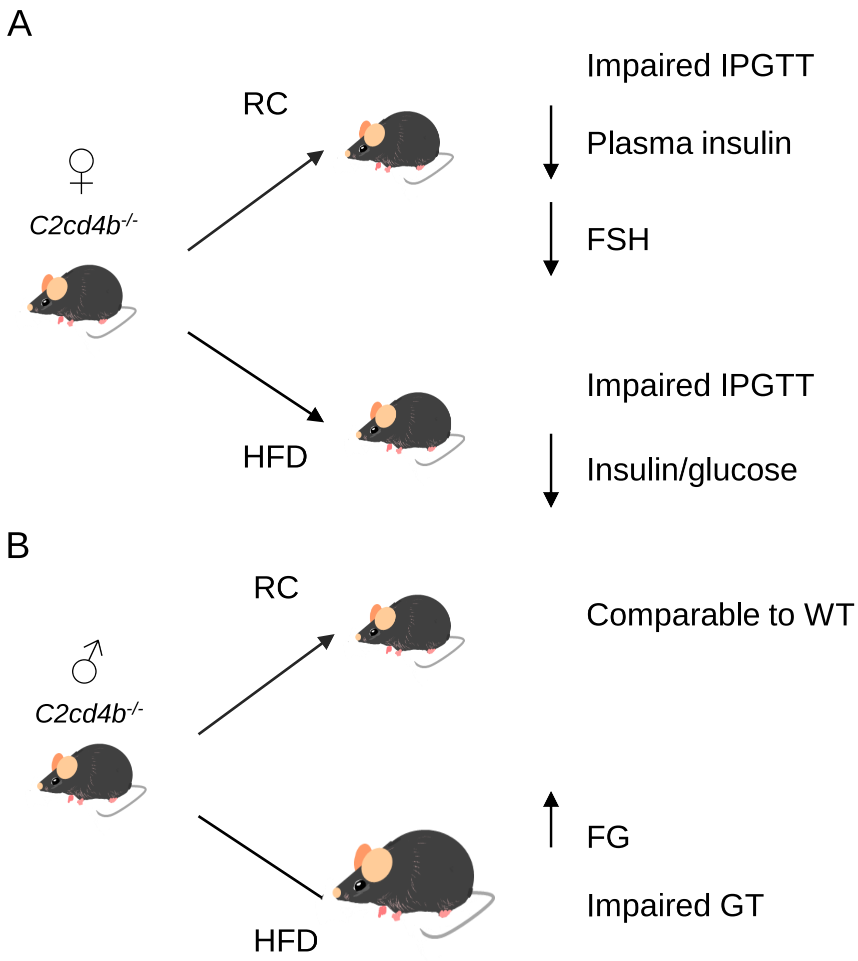


Supplementary Figure 17. Summary of the effect of deletion of *C2cd4b* on glucose homeostasis in mice. A. Female *C2cd4b* null mice showed impaired glucose tolerance on either RC or HFD, decreased plasma insulin and FSH levels when maintained on a RC, and a decreased plasma insulin to glucose ratio when maintained of HFD. B. *C2cd4b* null male mice showed comparable glucose homeostasis to WT littermates when maintained on RC. On HFD they showed glucose intolerance, increased lean body mass and elevated fasting glycemia. FG: fasting glucose, GT: glucose tolerance, WT: wild types.
